# A comprehensive survey of long-range tertiary interactions and motifs in non-coding RNA structures

**DOI:** 10.1101/2022.11.01.514747

**Authors:** Davyd R. Bohdan, Valeria V. Voronina, Janusz M. Bujnicki, Eugene F. Baulin

## Abstract

Understanding the 3D structure of RNA is key to understanding RNA function. RNA 3D structure is modular and can be seen as a composition of building blocks of various sizes called tertiary motifs. Currently, long-range motifs formed between distant loops and helical regions are largely less studied than the local motifs determined by the RNA secondary structure. We surveyed long-range tertiary interactions and motifs in a non-redundant set of non-coding RNA 3D structures. A new dataset of annotated LOng-RAnge RNA 3D modules (LORA) was built using an approach that does not rely on the automatic annotations of non-canonical interactions. An original algorithm, ARTEM, was developed for annotation-, sequence- and topology-independent superposition of two arbitrary RNA 3D modules. The proposed methods allowed us to identify and describe the most common long-range RNA tertiary motifs. Three basic interaction types were identified to be recurrent in the long-range RNA 3D modules: ribose-ribose interactions, canonical Type I and Type II A-minor interactions, and previously undescribed staple interactions. These three interaction types were found to be different building blocks of the same complex staple motifs common to non-coding RNA 3D structures.

## INTRODUCTION

Non-coding RNAs play important roles in many cellular processes [1]. Like that of proteins, the functions of non-coding RNAs are largely determined by their spatial structure [2]. RNA 3D structure is modular and can be seen as a composition of building blocks of various sizes called tertiary motifs [3,4]. Such recurrent motifs are often functionally important and structurally conserved [5–8]. Generally, all the motifs are defined based on hydrogen bonding interactions [6, 9–12], hydrophobic stacking interactions [13–15], or both of them [16–19]. The structural context matters for RNA tertiary motifs formation and can be leveraged for the prediction of their location [20,21].

In the context of RNA secondary structure, tertiary motifs can be divided into two groups - local motifs and long-range motifs. Local motifs include interactions within a secondary structure element or a number of adjacent elements, e.g., a GNRA tetraloop motif is a 4-nucleotide hairpin of a particular sequence and geometric configuration [16], a kink-turn motif is a sharp turn in phosphodiester backbone formed by two stems and an internal loop between them [17], a coaxial helix motif is a coaxial arrangement of two stems that usually share a common multiple junction [22]. Therefore, local motifs are subject to a natural comprehensive classification based on five standard secondary structure elements: a hairpin, a bulge, an internal loop, a multiple junction, and a stem. Currently, the classification is utilized in a great number of databases that list local tertiary motifs annotated in known RNA 3D structures [23–28].

In turn, long-range motifs are formed by interactions between distant secondary structure elements and, at the moment, lack a comprehensive classification. A few specific cases of such motifs were characterized, e.g., GAAA/11nt motif formed by a hairpin of GAAA sequence and an 11-nucleotide internal loop [29], D-loop/T-loop - an intricate network of interactions between two hairpins [7, 30–32], and A-patch - a continuous stack of A-minor interactions [6]. A number of works describe motifs that contain particular interactions that can be either local or long-range, e.g., A-minor motifs [20], ribose zippers [9], ribose-phosphate zippers [33], and base-intercalated or base-wedged elements (bie/bwe) [13]. The databases that include long-range motifs are often focused on these described types [34,35]. The majority of existing approaches consider only motifs formed by base-pairing interactions and completely ignore other hydrogen bonds or stacking interactions [21, 35, 36]. Therefore, the annotation of the motifs relies on the automatic annotation of the base-pairing interactions, which is usually performed with one of the five standard tools [37–41]. Furthermore, the pseudoknots are often excluded from the consideration by means of a popular procedure named “pseudoknot removal” [35].

Unlike local motifs that represent separate secondary structure elements and normally do not intersect with each other, long-range motifs often share nucleotides, e.g., GAAA/11nt motifs always include A-minor interactions [20], and A-minor interactions very often intersect with ribose zippers [30]. At the moment, few studies analyzed this kind of inter-motif relationships [20,30].

Currently, there exist a wide number of different tools for the pairwise superposition of RNA 3D structures, which can be divided into two main groups based on the specific purpose they were developed to fulfill. The majority of the tools focus on producing a structure-based RNA sequence alignment [42–49] and therefore usually require single RNA chains as input [43, 44, 47]. Another large group of tools is focused on local tertiary motifs superposition and search [50–53]. Hence, these tools rely on some of the structural features of the local motifs, e.g., on sequence conservation, annotation of canonical and non-canonical interactions, and backbone topology. Among the least constrained of the currently available tools which can be considered of general use are SETTER [54] and CLICK [55]. However, SETTER relies on RNA secondary structure elements, and CLICK is designed to produce RNA sequence alignment. To the best of our knowledge, up to the present moment, there was no tool specifically designed for the comparison of two arbitrary RNA 3D modules, able to find local and global structural similarities simultaneously, and independent of RNA sequence, RNA backbone topology, and annotation of any interactions.

In this work, we performed a comprehensive survey of long-range RNA tertiary interactions and motifs. We selected a dataset of long-range nucleotide doublets using a capacious < 6.0 Å threshold and the generalized RNA secondary structure description model that allows pseudoknots [20]. The doublets were grouped into long-range RNA 3D modules using a newly developed approach that does not rely on the annotation of any interaction types, only on spatial proximity. The assembled 3D modules were then used to derive motif instances and identify common long-range tertiary motifs via a new approach to the pairwise superposition of RNA 3D modules. The results confirm that A-minor-like motifs are the most populated types of long-range contacts in non-coding RNA 3D structures, but also discover a number of previously undescribed interactions. Furthermore, the results show large disagreements between the standard automatic RNA structure annotation tools.

The proposed original algorithms make it possible to perform sequence-, topology- and annotation-independent description, analysis, and superposition of the RNA structural modules, and are not limited to long-range motifs. We believe the proposed methods will have a great impact on comparative analyses of both RNA structures and RNA-containing complexes.

## RESULTS

### 1. Automatic annotation tools should not be used for assembling long-range RNA 3D modules

In this work we defined a long-range nucleotide doublet as two ribonucleotides from distant RNA secondary structure elements, stems or loops, with any non-hydrogen atoms within 6.0 Å of each other, see the Materials and Methods section. The collected dataset included 17,272 long-range doublets (Table S1) formed by 11,204 nucleotides from 70 RNA chains of the non-redundant set of RNA structures (Table S2). 16,078 (93%) doublets were formed by 10,714 (95.6%) nucleotides of a standard residue type (32.3% G, 26.3% A, 23.4% C, 18% U) that were assigned with a particular conformation (anti/syn ~C3’-endo/~C2’-endo) using DSSR [38]. The dataset included 221 (2%) modified residues. 1554 (9%) of the long-range doublets are intermolecular doublets of nucleotides from different RNA chains forming a shared RNA secondary structure.

The doublet nucleotides are almost equally split between stems (48%) and loops (52%). However, more than 80% of the adenosines belong to loops, whereas 69% of the cytidines and 61% of the guanosines belong to stems, and uridines show the most equal split with 57% of loop residues. Pseudoknotted stems and loops compose only 16% of the doublet residues, with an almost two-fold difference between stems (5.5%) and loops (10.5%). Among the 84% of classical (pseudoknot-free) stems and loops the largest share belongs to stems (42%), followed by hairpins (18%), internal loops and bulges (15% combined), and multiple junctions (9%).

Stem-loop doublets form 55% of the dataset, and loop-loop and stem-stem doublets form 30% and 15% respectively. A(loop)-G(stem) and A(loop)-C(stem) are the most common long-range doublets representing together 20.3% of the dataset (11.8% + 8.5%). The third most common doublet is A(loop)-G(loop) (5.8%). The most frequent pairs of RNA secondary structure elements among the long-range doublets are stem-hairpin (16.8%), stem-internal loop (12.4%), stem-stem (11.7%), stem-multiple junction (7.6%), and hairpin-hairpin (5.4%).

Among the closest atoms of the long-range doublets, the most frequent are O2’ (20%) and OP1 (14.3%), followed by O4’ (8.8%). Among base-specific atoms, the most frequent are N2G (5.3%), C2A (3.9%), N1A (3.3%), and N6A (3.2%). The most frequent pairs of closest atoms in the long-range doublets are O2’-OP1 (7%) and O2’-O2’ (5.6%), and C2A-N2G (1.3%) is the most frequent base-specific pair of atoms.

The long-range doublets were annotated with the previously defined interaction and motif types using the five conventional automatic annotation tools: FR3D [37], DSSR [38], MC-Annotate [39], ClaRNA [40], and RNAView [41], see the Materials and Methods section.

DSSR annotated 3,244 (18.8%) doublets as non-pairing H-bonds (H-bonds that are not a part of canonical or non-canonical base pair) and 1,345 (7.8%) doublets as A-minor interactions. The five tools annotated on average 1,153 (6.7%) doublets as base pairs, ClaRNA being the most conservative tool (553 base pairs) and RNAView the least conservative (1,625 base pairs). Simultaneously, RNAView annotated as base stacking just 26 doublets, whereas FR3D annotated 819 (4.7%) doublets, and the other three tools on average annotated 446 doublets.

Each of the annotated interaction and motif types is dominated by loop adenosines composing from 25% (non-pairing H-bonds) to 56.3% (base-intercalated elements [13]) of the interacting residues. Among all the 11,204 nucleotides forming the long-range doublets loop adenosines compose 21%. Base stacking interactions are dominated with A(loop)-A(loop) doublets (17%-36%). Almost all of the other interaction and motif types are dominated by A(loop)-G(stem) doublets (12%-38%), whereas A(loop)-C(loop) doublets prevail in DSSR ribose-zippers (36%) and non-pairing H-bonds (13%).

Strikingly, among 2919 long-range doublets annotated as a base pair by at least one tool, only 13% are consensus five-tool annotations and almost 49% are single-tool annotations.

We further examined the base pair annotations separately for each Leontis-Westhof type [11]. The Leontis-Westhof type (LW type) is a three-letter code specifying the first letters of the two interacting base edges (*Watson-Crick/Hoogsteen/Sugar*) along with their relative configuration (*cis/trans*), e.g., *tHS* is an interaction between Hoogsteen edge of one base and Sugar edge of another base with the bases being in *trans* configuration. Among the base pairs of determined LW-type annotated in the long-range doublets, four types *cSW/tSW/tSS/cSS* compose from 60% to 83% depending on the tool, and types that involve a Hoogsteen edge compose just 7%-16%. Surprisingly, single-tool annotations dominate ten of the twelve possible LW types, composing more than 50% in eight of the ten. Two exceptions, *cWW* and *tWW* types are dominated by four-tool (50.6%) and five-tool (38.6%) annotations respectively.

We discovered 180 long-range doublets annotated with different base pair LW-types by different tools (Table S3, Figure 1). A single example of a doublet annotated with three different LW types was found (Figure 1D) and the other 179 doublets were annotated with two LW types. 148 (82%) of such doublets involve adenosines (66 A-G, 53 A-C, 17 A-A, 12 A-U). 143 (79%) are SS/SW disagreements (Figure 1A, B), and for 91 of them, among other tools, FR3D annotates an SS type and DSSR annotates an SW type.

**Figure 1.**
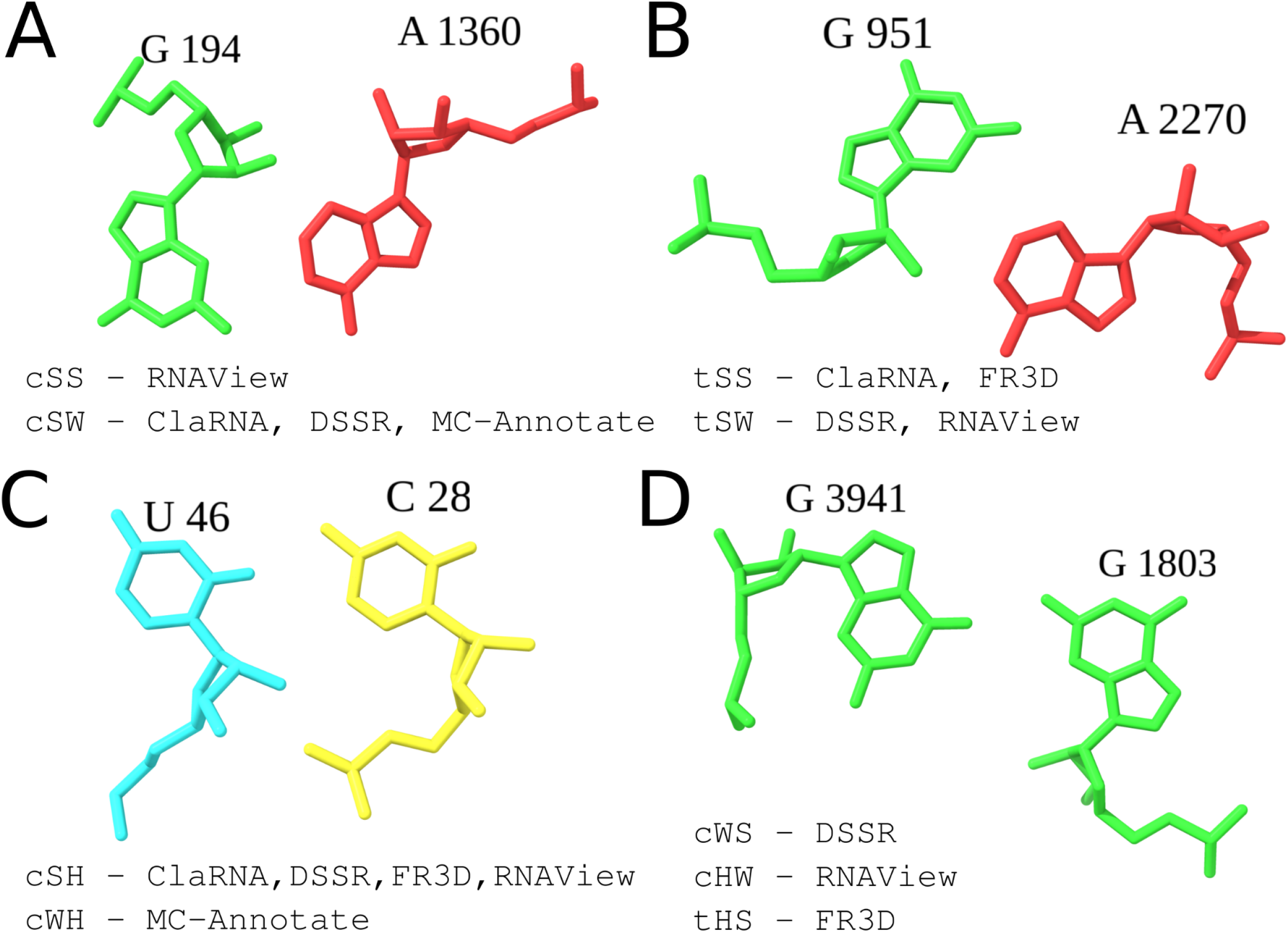
Examples of long-range nucleotide doublets annotated with different Leontis-Westhof base pair types. (A) PDB entry 7M4Y, chain A; (B) PDB entry 7M4Y, chain A; (C) PDB entry 7DVQ, chain F; (D) PDB entry 7O7Y, chain B5.

Notably, *cSS/cSW/tSS/tSW* base pairs are common for ubiquitous A-minor interactions, hence their annotation is of the highest importance for the assembly of long-range RNA 3D modules.

Furthermore, 10,379 (60%) of the long-range doublets were not annotated with any common interaction type using any of the tools. It means they would be totally missed in the annotation-based assembly of RNA 3D modules, and that includes 2,095 doublets (12%) that involve at least one pair of non-hydrogen atoms under 4.0 Å.

### 2. An overview of the long-range RNA 3D modules dataset

The dataset of 17,272 long-range doublets was used to assemble the LORA (LOng-RAnge) dataset of 1,264 long-range RNA 3D modules. For that, we developed an original approach that does not rely on the automatic annotations of non-canonical interactions, see the Materials and Methods section.

The LORA dataset included 389 modules of size two (i.e., composed of two residues), 183 modules of size three, 686 modules of sizes from 4 to 81 (Figure S1), and six modules from ribosomal RNAs (rRNAs) larger than 100 residues in size. A unique identifier made of a PDB entry and an ordinal number was assigned to each module, e.g., *“7o7y_7”* (Table S1).

To assess the suitability of the module assembly we examined instances of the well-known “named” long-range RNA tertiary motifs (Figure 2). Module *6ugg_0* (15 residues) is a canonical instance of the D-loop/T-loop interaction motif from tRNA (Figure 2A). Module *6ugg_0* includes two long-range base pairs *G17-U55(tWS*) and *G18-C56(cWW), both* annotated with all five tools, and an independent long-range base stacking *G14-G59* annotated with all the tools except RNAView.

**Figure 2.**
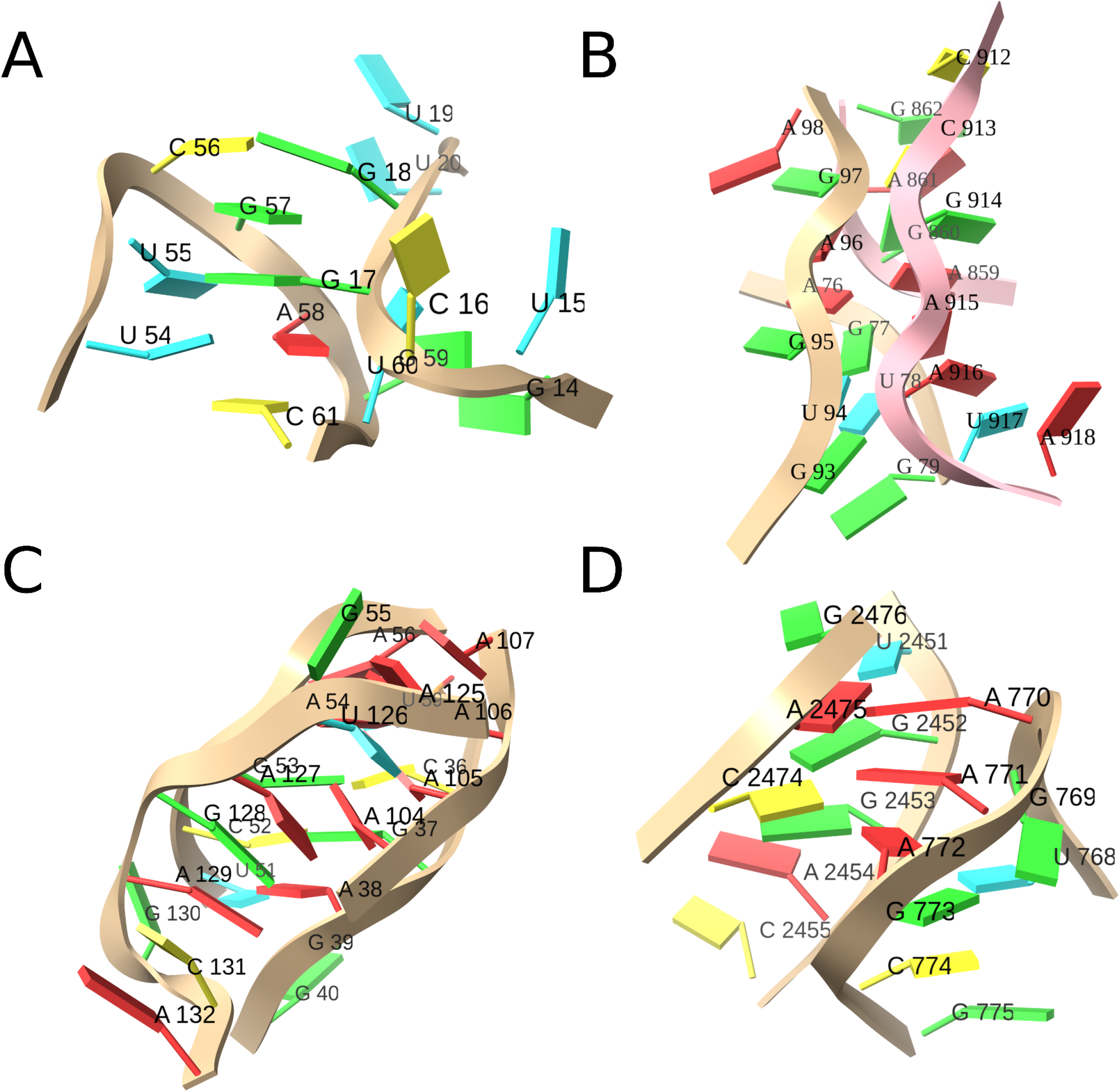
Examples of the assembled long-range RNA 3D modules belonging to “named” tertiary motifs. (A) D-loop/T-loop interaction motif, module 6ugg_0, PDB entry 6UGG, chain B; (B) Double symmetric A-patch, module 7m4y_45, PDB entry 7M4Y, chains A (light pink) and B (beige); (C) A-patch, module 2z75_1, PDB entry 2Z75, chain B; (D) GNRA-tetraloop/tetraloop-receptor, module 6th6_41, PDB entry 6TH6, chain BA.

Module *7m4y_45* (21 residues) is a double symmetrical A-patch formed between 23S and 5S rRNAs (Figure 2B). The A-patch is the only conserved tertiary interface between 23S and 5S rRNAs [6], and it is conserved throughout all the kingdoms of life. Our dataset includes three instances of this A-patch - modules *7m4y_45, 6th6_57*, and *7o7y_11*. In module *7m4y_45* 5S rRNA purines *A76, A96*, and *G97* interact with the minor groove of 23S rRNA near base pair *G860-C913(cWW*), and in turn, 23S rRNA purines *A859, A915*, and *A916* interact with the minor groove of 5S rRNA near base pair *G77-U94(cWW*). Here, DSSR and RNAView annotate the only long-range base pair *G77-A915(tSS*) and DSSR annotates zero long-range A-minors. In turn, FR3D annotates three base pairs of “near” geometry (i.e., interactions close to a base pair but not meeting the strict criteria): *G77-A859(ncSS*), *A96-G860(ntSS*), and *G77-A915(ntSS*), MC-Annotate annotates four “pairings” between an O2’ atom and a Sugar edge (*G77-A859, A76-G860, U78-A916*, and *G93-A916*), and ClaRNA annotates *G77-A915(tSS*) base pair with score 0.28. Hence, no automatic annotation tool identified the entire A-patch involving six purine bases.

Module *2z75_1* (24 residues) is an A-patch from glmS ribozyme, which involves a stack of six purine bases (*A129, G128, A127, A104, A105, A106*) interacting with the minor groove of canonical base pairs *A38-U51, G37-C52, C36-G53*, and *U59-A54* (Figure 2C). Here, DSSR annotates three long-range base pairs, but simultaneously it annotates five A-minors involving five of the six purine bases, except *A106*. FR3D annotates five long-range base pairs of “near” geometry which involve five of the six purine bases, except *A129*. MC-Annotate too annotates long-range interactions involving five of the six purines but not *A129*. ClaRNA annotates three long-range base pairs with a score above the threshold, and three more base pairs with scores below the threshold. RNAView is the only tool able to annotate here long-range base pairs involving all six purine bases.

Module 6th6_41 (16 residues) is an instance of the GNRA-tetraloop/tetraloop receptor interaction motif (Figure 2D). Here a GAAA tetraloop (*G769*, *A770, A771, A772*) interacts with the minor groove of canonical base pairs *U2451-A2475* and *G2452-C2474*. Here all the tools annotate the long-range interaction involving *A770*, but RNAView and MC-Annotate fail to annotate the long-range interaction involving *A771*.

Overall, none of the five automatic annotation tools is able to fully annotate more than two of the four described RNA 3D modules, and with all of the tools simultaneously still it would be possible to annotate only three of the four modules.

### 3. Exploring sequence-, topology-, and annotation-independent structural similarities of D-loop/T-loop-like motifs

To compare the assembled long-range RNA 3D modules we developed a new algorithm for pairwise structural superposition, ARTEM (Aligning Rna TErtiary Motifs). The ARTEM algorithm is designed to find structural similarities between two arbitrary RNA 3D modules of possibly different sizes and without a priori knowledge of the nucleotide matchings between the modules, see the Materials and Methods section.

To demonstrate the capabilities of the ARTEM algorithm we performed a search for RNA 3D modules structurally similar to the D-loop/T-loop interaction motif from tRNA represented in the LORA dataset with module *6ugg_0*, (Figure 3A, C). We searched for the similarities with *RMSD ≤ 3.0 Å, SIZE ≥ 7 residues*, and *RMSD/SIZE ratio ≤ 0.25* that involve at least one long-range nucleotide doublet within the superimposed subset of residues. These thresholds were chosen to filter out non-relevant hits, e.g., two- or three-residue hits with RMSD close to zero, or hits with no long-range doublets.

**Figure 3.**
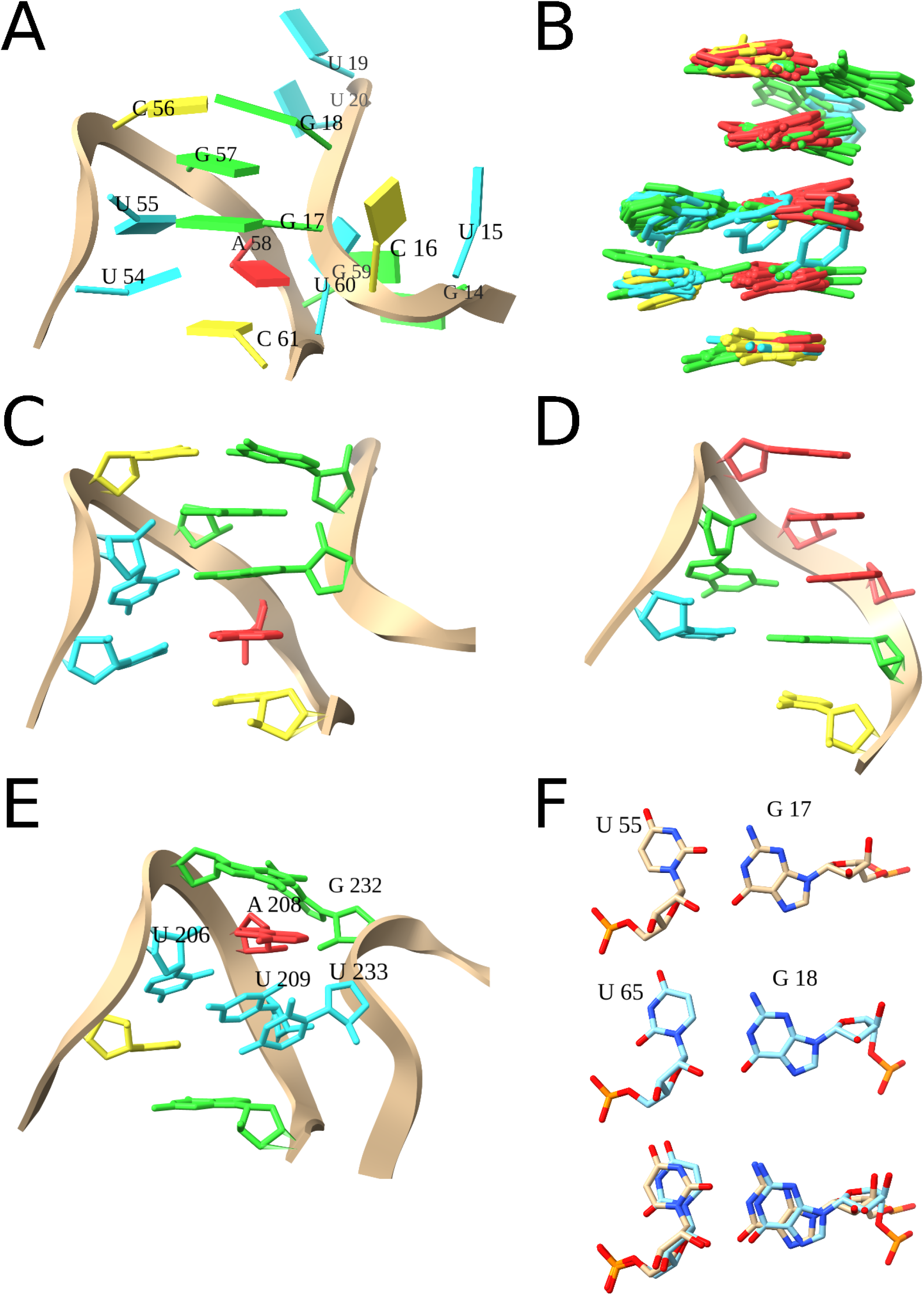
D-loop/T-loop-like RNA tertiary motifs. (A) Reference D-loop/T-loop motif from tRNA (module 6ugg_0). (B) Superposition of the 32 motifs structurally similar to the D-loop/T-loop motif. (C) The structurally conserved part of the reference 6ugg_0 module. (D) The GAAA-tetraloop (module 6th6_41). (E) D-loop/T-loop-like motif from eukaryotic LSU rRNA (module 7o7y_58). (F) tSW U55(anti)-G17(anti) base pair from module 6ugg_0 (top), tHW U65(syn)-G18(anti) base pair from module 3q1q_0 (middle), and their superposition (bottom). In panels A-E the standard residue-coloring scheme is used with adenosines in red, uridines in blue, cytosines in yellow, and guanosines in green.

Overall we found 32 hits (Table S4, Figure 3B). The D-loop/T-loop-like motifs were found in tRNA (modules *3rg5_0, 3q1q_0*), Y RNA (*6cu1_0*), virus tRNA-like UTR (*6mj0_0*), Hatchet ribozyme (*6jq5_0*), RNase P (*2a64_0*), several riboswitches (e.g., *4frn_1* and *3f2x_1*), and multiple hits were found in archaeal, bacterial, and eukaryotic rRNAs.

Three of the found motifs are tetraloops (GAAA in *6th6_41*, GCAA in *7m4y_3*, and UAAC in *3q1q_3*) forming a nested non-canonical base pair in place of the long-range base pair *U55-G17(tSW*) in the D-loop/T-loop motif (Figure 3D). The found tetraloops show that ARTEM can find similar RNA base arrangements in a topology-independent manner. Also, out of 29 non-tetraloop hits, only 14 are hairpin-hairpin interfaces as is the reference module *6ugg_0*.

Module *7o7y_58* is the only one that has two pyrimidines matching the highly conserved *U54-A58(tWH*) base pair in *6ugg_0*, and their geometry differs from a common arrangement of the D-loop/T-loop motif (Figure 3E). Here the *U233* residue stacks under *U209* instead of stacking between *U209* and *A208*, and this arrangement precludes *U233* from forming a long-range base pair with *U206* that would match the *U55-G17(tSW*) base pair in *6ugg_0*. Thereby, ARTEM found a motif that is similar to the D-loop/T-loop but has an alternative arrangement of the residues, which would not be possible using the base pair annotation-based search.

In place of the long-range *U55-G17(tSW*) base pair in *6ugg_0*, module *3q1q_0* has an unusual *U65(syn)-G18(anti)(tHW*) base pair that was not annotated by any automatic annotation tool (Figure 3F). The superposition of the two base pairs suggests that it may be a case of modeling error and the residue *U65* was wrongly put into *syn* conformation. Hence, ARTEM can be of service in resolving the conserved structural motifs in RNA structure modeling.

### 4. Ribose-ribose interactions, canonical A-minor interactions, and previously undescribed staple interactions form the most common long-range RNA tertiary motifs

To identify commonly occurring long-range RNA tertiary motifs we applied the developed ARTEM algorithm to the long-range RNA 3D modules of the LORA dataset. First, we identified more than a million motif instances and then clustered them to derive the most populated motif types. We assigned unique identifiers to motifs and motif instances, e.g., motif instance *“7m4y_128_455”* of motif *“555_8_0.36_7m4y_128_455”*, see the Materials and Methods section.

Overall, we identified 1,156,112 motif instances of sizes from 2 to 375 (Figure S2). The further analysis was limited to motif sizes from 2 to 30, as only a negligible quantity (3,149) of instances were larger than 30 residues in size. After applying an original heuristic clustering procedure we identified 2,733 motifs of sizes from 2 to 30. Among 424 motifs populated with at least 100 instances, only motifs of sizes from 3 to 11 were represented (Figure S3). All the 424 motifs with at least 100 instances are made of various interactions of the minor groove of a helical region with up to three residues. Such interactions are known to be by far the most common long-range interactions in non-coding RNA 3D structures [6,56].

We then further visually examined the top 100 most frequent motifs and grouped them based on their hydrogen bond networks (Table 1, Table S5). The most populated group included 26 motifs with a single ribose-ribose O2’-O2’ interaction. The second most populated group included 22 motifs incorporating a canonical Type I A-minor interaction. During the visual examination, we noted a significant number of motifs with one or two residues interacting simultaneously with both helical strands at different base pair layers (Figure 4). Overall we counted 32 such motifs which we called *“staples”*. In every staple, at least one O2’/O4’ atom of each helical strand is involved in an interaction with the third strand, the *staple strand*. If the interactions involved only ribose atoms of a staple residue, we included “N” in its name to emphasize the lack of base specificity, e.g., NN3-staple is an interaction of a helix region with two non-helix riboses spanning three base pair layers of the helix (Figure 4B). According to the proposed naming, Type I A-minor is usually a case of A1-staple, as here interactions are limited to a single base pair layer and are highly specific to the adenine base. However, in the context of the symmetric Type I/I A-minor motif involving a cross-strand stack of adenosines, Type I A-minor interaction can be an A2-staple, see e.g., motif *555_8_0.36_7m4y_128_455*.

**Figure 4.**
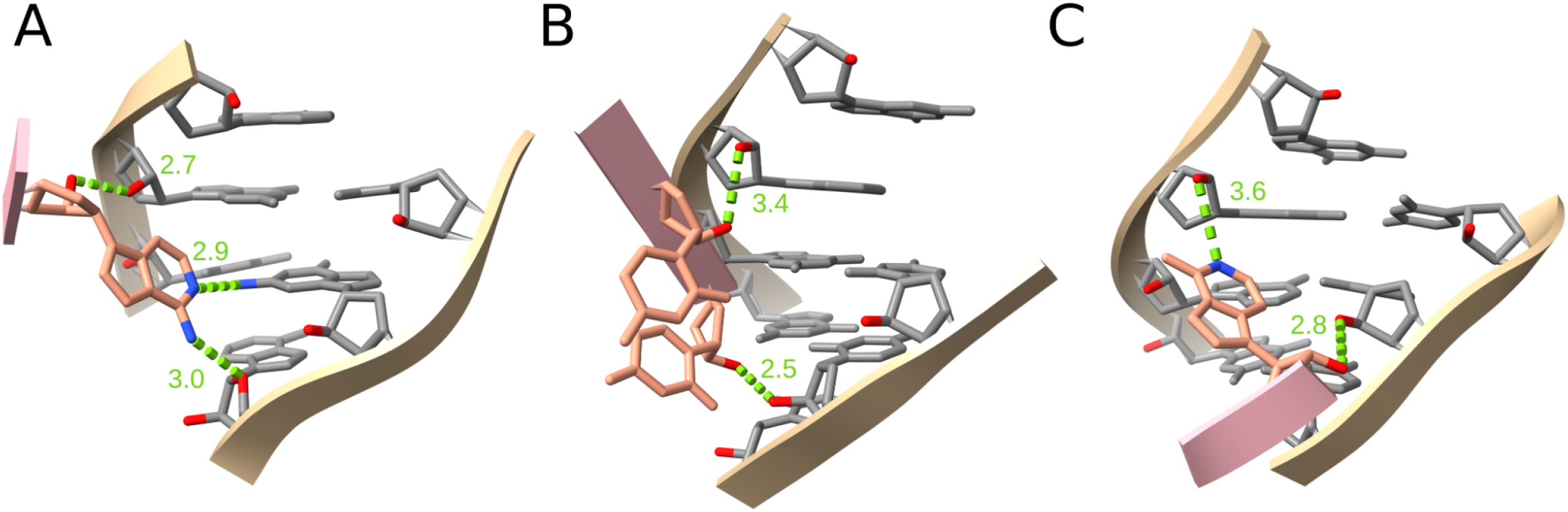
Staple interaction examples. (A) A3-staple, motif 539_7_0.26_2z75_1_259, (B) NN3-staple, motif 482_9_1.00_7o7y_48_1140, (C) Type II A-minor (A2-staple), motif 450_8_0.40_3p49_0_706. The staple strands are shown in pink and salmon. Interacting N and O atoms are shown in blue and red respectively. O2’ atoms are shown in red to designate the minor groove. H-bonds are shown in green along with their distance in angstroms.

**Table 1.**
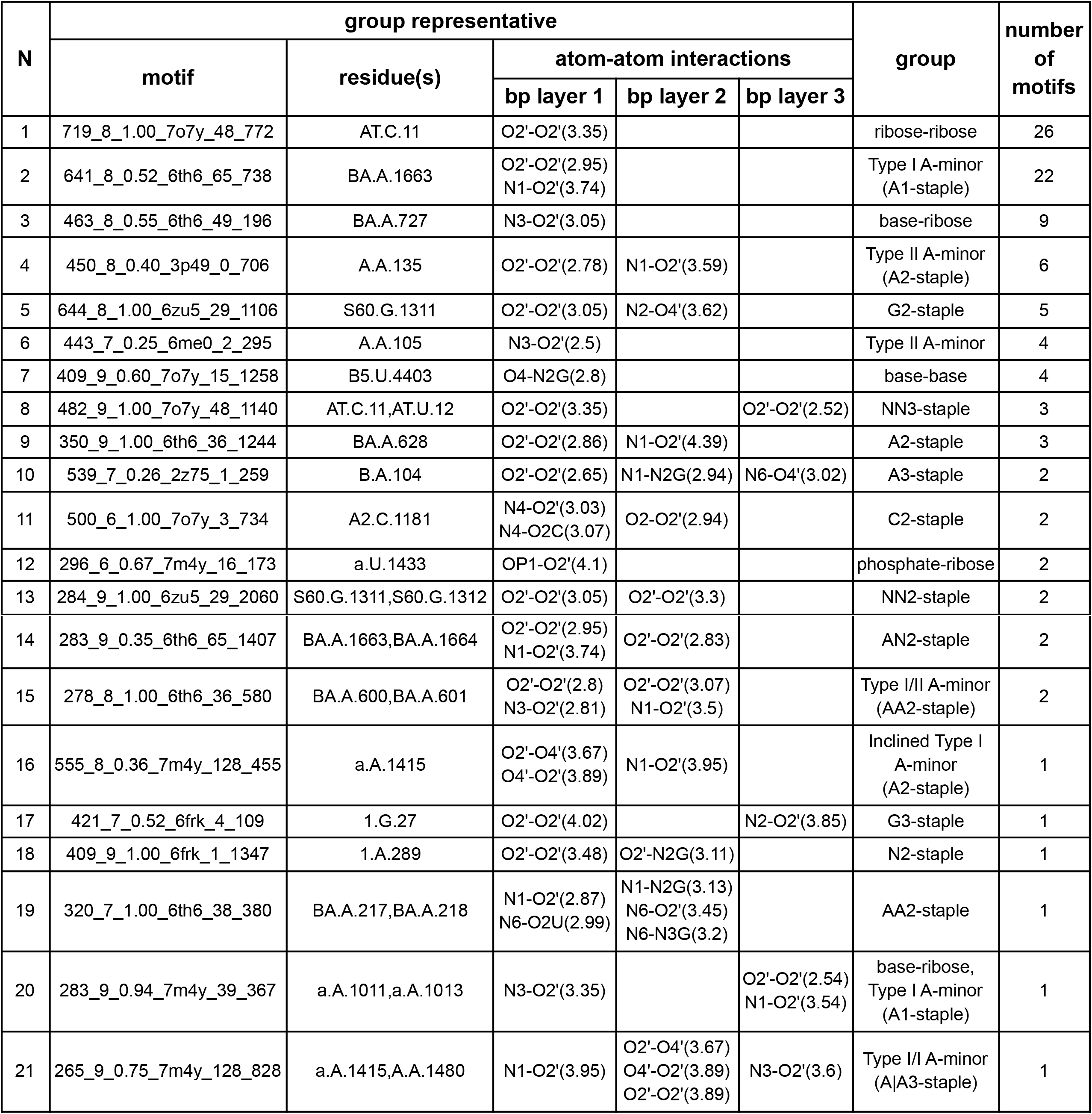
The top 100 most frequent motifs form 21 groups based on their interaction networks.

To better understand the nature of the observed motifs we further visually examined their source LORA modules to see the structural context of the motifs (Table S5). Surprisingly, non-helical residues of the 98 out of 100 most frequent motifs belonged to a larger staple strand or two staple strands with a cross-strand stack. From the 50 structural contexts of the 98 motifs, we counted 15 AAN3-staples, 12 NA|AAN4-staples (here pipe represents a cross-strand stack), and 9 NN3-staples (Figure 5, Figure 4B).

**Figure 5.**
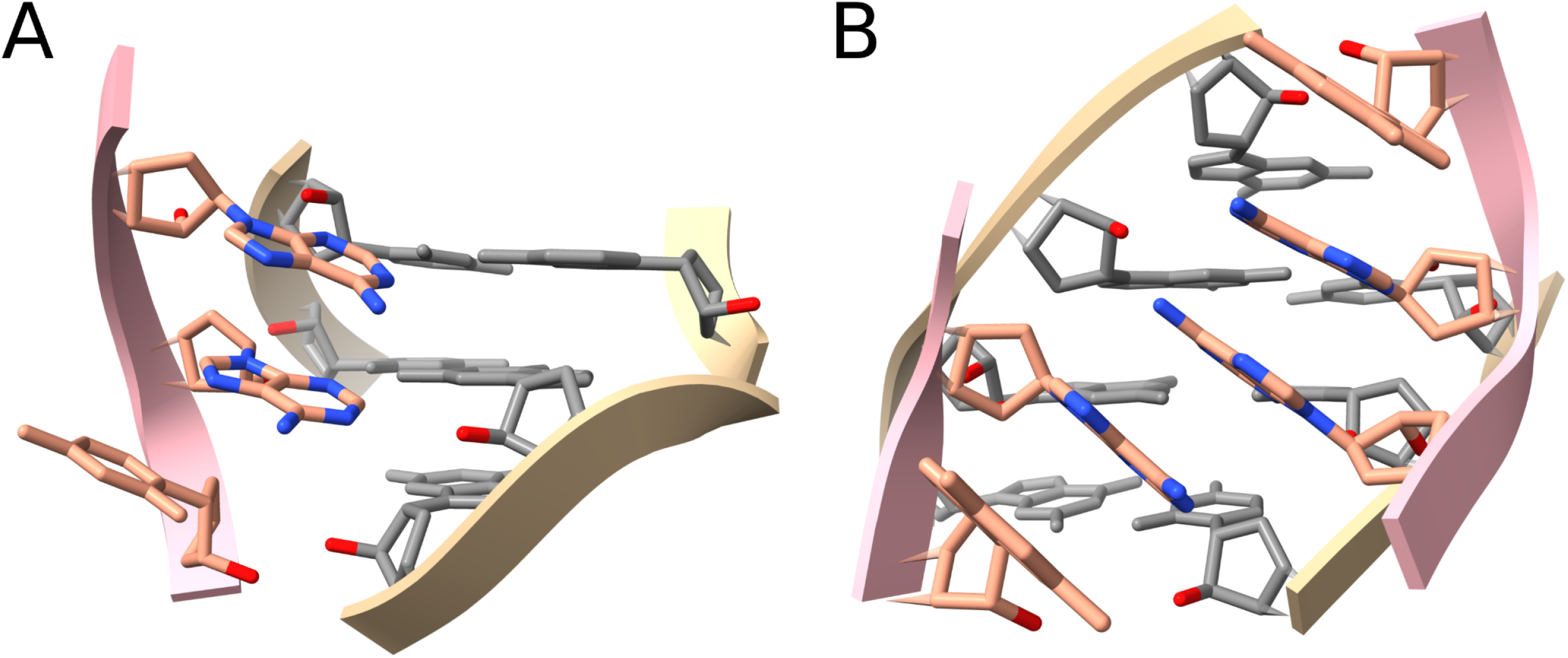
Complex staple motif examples. (A) AAN3-staple motif from module 7ez2_1 (PDB entry 7EZ2, chain N, residues A152, A153, C154), (B) NA\AAN4-staple motif from module 6th6_36 (PDB entry 6TH6, chain BA, residues G629, A628, A600, A601, G602). The staple strands are shown in pink and salmon. N atoms of the staple adenosines are shown in blue. O2’ atoms are shown in red to designate the minor groove.

To confirm that staples are not only found in rRNAs that dominate the LORA dataset, we examined the instances of motif *539_7_0.26_2z75_1_259* from the A3-staple group and found the group representatives from seven different RNA molecule classes (Table 2). In two out of seven representatives the staple residue is guanosine. Although, due to the limitations of the used clustering approach we observed that the majority of motif *539_7_0.26_2z75_1_259* instances belong to the A2-staple group rather than the A3-staple group, which turned out to be rare.

**Table 2.** A3-staple group motif 539_7_0.26_2z75_1_259 representatives from different non-coding RNA molecules.

| PDB entry | RNA molecule | Motif instance | Staple residue | Layer-1 | Layer-2 | Layer-3 |
| --- | --- | --- | --- | --- | --- | --- |
| 2Z75 | glmS ribozyme | 2z75_1_259 | B.A.104 | O2'-O2'(2.65) | N1-N2G(2.94) | N6-O4'(3.02) |
| 6TH6 | Archaeal SSU rRNA | 6th6_48_678 | Aa.A.1239 | O2'-O2'(3.48) | N1-N2G(3.04) | N6-O4'(3.55) |
| 6TH6 | Archaeal LSU rRNA | 6th6_27_968 | BA.A.1716. | O2'-O2'(3.1) | N1-N2G(2.89) | N6-O4'(3.34) |
| 4WFL | Bacterial large SRP RNA | 4wfl_0_354 | A.G.76. | O2'-O2'(2.78) | - | N2-O4'(3.8) |
| 5T83 | Guanidine-I riboswitch | 5t83_0_210 | A.A.82. | O4'-O2'(3.31)<br>O2'-O4'(3.29) | N3-O2'(2.73) | N6-O2'(3.16) |
| 7D1A | Group II catalytic intron | 7d1a_4_746 | A.G.501. | O4'-O2'(2.91) | N1-O2'(2.3) | N2-O4'(2.71) |
| 7EZ2 | Group I catalytic intron | 7ez2_0_400 | N.A.115. | O4'-O2'(3.09) | O2'-O2'(3.78) | C2-O2'(3.75) |

During the analysis of the most frequent motifs and their structural context, we also noted a number of interesting topology-independent structural similarities between instances of the same motif (Figure 6). Here, on panel A, motif instance *3ndb_0_941* incorporates a Type I/II A-minor motif with a canonical base pair and discontinuous non-canonical *A-U(tHW*) base pair, whereas on panel B motif instance *6th6_18_261* forms a Type I/II A-minor motif with two canonical base pairs but the highest base pair layer is formed with the staple strand, not the helical strand. Another case is shown in panels C and D, where adenosine forming a canonical Type I A-minor interaction in *6th6_65_738* matches a C-minor-forming cytidine consecutive to one of the canonical base pair residues in *6zu5_5_3291*. The two examples demonstrate that in RNA structures specific base arrangements can be preserved even within different backbone topology contexts.

**Figure 6.**
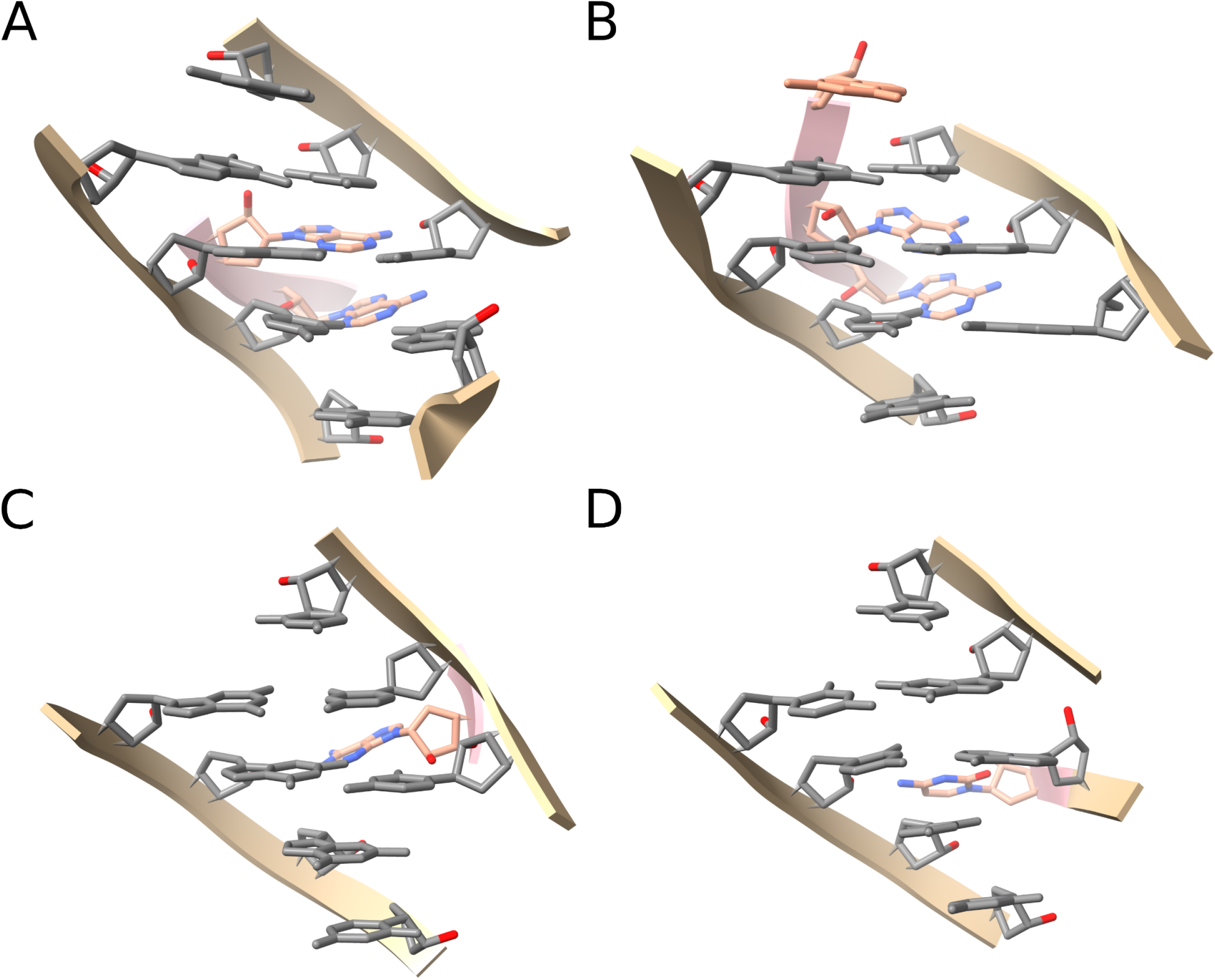
Topology-independent structural similarities between motif instances. (A) Instance 3ndb_0_941 and (B) instance 6th6_18_261 from motif 241_10_1.00_7o7y_48_1570; (C) Instance 6th6_65_738 and (D) instance 6zu5_5_3291 from motif 641_8_0.52_6th6_65_738. The staple strands are shown in pink and salmon. N and O atoms of the staple bases are shown in blue and red. O2’ atoms are shown in red to designate the minor groove.

## DISCUSSION

In this work, we performed a survey of long-range tertiary interactions and motifs in a non-redundant set of non-coding RNA 3D structures. First, we identified long-range nucleotide doublets based on the capacious 6.0 Å threshold and the annotation of RNA secondary structure. Then we grouped the 17,272 long-range doublets into the 3D modules using an original approach based purely on the proximity of the doublets. The assembled 1,264 long-range modules composed the LORA dataset. And finally, we applied ARTEM, a novel algorithm for RNA tertiary motifs pairwise superposition, to derive more than a million long-range tertiary motif instances from the 3D modules of the LORA dataset, which were later clustered into tertiary motifs. The 100 most common long-range tertiary motifs were then analyzed in detail.

We found large disagreements between the five tools commonly used for the automatic annotation of non-canonical interactions in RNA structures, in particular for relatively uncommon interactions. Using the well-known “named” RNA tertiary motif examples we showed the high quality of the assembled long-range RNA 3D modules, which could not be achieved based on the automatic annotation of the involved interactions. Searching for D-loop/T-loop-like motifs we demonstrated the capability of the developed ARTEM algorithm to find structural similarities with different backbone topology or interaction network variations and even find potential modeling errors in place of the supposedly highly conserved base pairs. Along with the well-known Type I and Type II A-minor interactions we identified a multitude of ribose-ribose interactions and previously undescribed interactions of one or two residues with the minor groove of a helical region simultaneously with both helical strands and at different base pair layers, which we called staples. Knowing the structural context of the motifs, we found these three types of interactions (canonical A-minors, ribose-ribose interactions, and staples) to be different building blocks of the same complex staple motifs common to non-coding RNA 3D structures.

In this work, we used a non-redundant set of RNA structures with one structure per non-coding RNA family from Rfam [57] rather than the more commonly used representative set of RNA structures with one structure per a pair of a source organism and an RNA molecule type [58]. This was done for two purposes, first, to automatically eliminate all synthetic and short RNA strands not linked to any Rfam family, and second, to fight the “ribosomal RNA curse”. Thus, among the representative structures linked to Rfam families the share of non-rRNA chains is 71.5% and the share of non-rRNA residues is only 18.26%, whereas in the used non-redundant subset the share of non-rRNA chains is 88.7% and the share of non-rRNA residues is 41.62% (Table S2).

Compared to the existing databases of long-range RNA tertiary motifs [35] the LORA dataset has the advantage of including the entire structural context of the long-range interactions, as it does not rely on automatic annotation of non-canonical interactions. LORA only relies on the RNA secondary structure annotation used to separate the long-range doublets from the local ones, but as it was shown previously [40] and also shown in this work, the automatic annotation tools agree well in the annotation of canonical cWW base pairs, i.e., base pairs forming the RNA secondary structure.

The most important limitation of the ARTEM algorithm is simultaneously its most important advantage. It produces structural superpositions based purely on RMSD in annotation-, sequence- and topology-independent manner. Accordingly, ARTEM does not distinguish between e.g., Type I and Type II A-minor interactions, especially in large structural contexts, as these two interactions are almost identical in their base arrangement and differ only in a small shift of the adenosine residue. Thus, in the analyses performed in this work the motif with the A3-staple group representative included many instances of A2-staples. Therefore, ARTEM identifies matches between spatially similar motifs that differ in topology, yet simultaneously it almost completely eliminates false negatives. Searching with ARTEM for a given reference motif in a given query RNA structure with no RMSD or SIZE thresholds it’s impossible to miss any of the motif occurrences unless an occurrence has zero residues that perfectly align with their reference counterpart, but in such a case it’s unclear why the occurrence is treated as the motif occurrence in the first place. E.g., ARTEM won’t identify a stack of two cSS base pairs as a motif structurally similar to a stack of two cWW base pairs.

The output of the ARTEM algorithm can be subject to additional filtering based on any required sequence specificity, interaction types, or backbone topology. Therefore, since ARTEM can be used for two arbitrary RNA structures or structural fragments, we believe the algorithm will find wide use in many applications. Furthermore, ARTEM can easily process any modified residues [59] provided their 5-atom representations are specified by the user.

In this work, we found three basic interaction types to be recurrent in the long-range RNA 3D modules: ribose-ribose interactions, canonical A-minor interactions, and staples. Among these three only the canonical A-minor interactions were comprehensively studied to the moment. We believe the other two were largely understudied due to a general approach bias toward base pair annotations, as e.g., the authors in [60] wrote: *“Further, enumerating a single feature such as a base-pair has a limited ability to describe the variety of structure seen in RNA; any non-paired structural elements, of which there are many, would fail to be described”*. Ribose-ribose interactions are too simplistic and not base-specific if considered in isolation. Staples by their nature are non-planar and cannot be decomposed into a set of base pairs. And only the canonical A-minor interactions can be naturally decomposed into a number of *cSS/tSS/cSW/tSW* base pairs.

We observed and demonstrated a number of topology-independent structural similarities, e.g., between a long-range D-loop/T-loop interaction motif and a GAAA-tetraloop hairpin. Previously, similar cases were described e.g., in [20] between an internal loop forming the across-bulged motif and the long-range GAAA/11nt interaction motif. The possibility of the kink-turn motif usually being an internal loop to occur in the context of a multiple junction is another example [61]. Taking this together we observe a strong ability of RNA structures to preserve specific base arrangements within different backbone topology contexts. Such topological differences between the instances of a conserved structural RNA motif can be relics of RNA backbone rearrangements, see e.g., [62]. Thus, we believe the scope of future RNA structural studies should not be limited to either only local or only long-range motifs but as well include comparisons between the two classes.

Summarizing, we performed a comprehensive analysis of the long-range tertiary interactions and motifs in non-coding RNA 3D structures. A new dataset of annotated long-range RNA 3D modules was built using the original approach which does not rely on the automatic annotations of non-canonical interactions. A new algorithm was developed for annotation-, sequence- and topology-independent superposition of two arbitrary RNA 3D modules. The proposed methods were used to identify and describe the recurrent long-range RNA tertiary motifs. We believe the results obtained in this work as well as the proposed methods will have a great impact in the field of RNA structural studies, especially in comparative analyses of RNA structures and RNA-containing complexes.

## MATERIALS AND METHODS

### 1. Dataset of long-range nucleotide doublets

To prepare a dataset of long-range nucleotide doublets from experimentally determined RNA 3D structures we used a non-redundant set of RNA structures from the Protein data bank (PDB, [63]). First, we downloaded the commonly used representative set of RNA structures from [58] (version 3.215 with 4.0 Å resolution cutoff). We annotated the representative RNA chains with the non-coding RNA family identifiers from Rfam [57] taking the family with the lowest E-value in case of multiple hits per RNA chain. Then, to construct a non-redundant set of RNA structures, we manually selected one RNA chain per family based on three features: resolution, number of resolved residues, and number of base pairs of any type, according to the annotation using the DSSR tool [38]. By default, we selected the chains with the best resolution, but in several cases, we did otherwise, e.g., when the best-resolution chain was a short fragment of the complete molecule, or when the second-best resolution chain was significantly longer and the difference in resolution was negligible. The resulting set included the structures of 97 non-coding RNA families from 89 PDB entries (Table S2).

To annotate RNA secondary structure elements and nucleotide doublets we used the urslib2 python library [20], which relies on DSSR annotations of canonical base pairs. We defined a nucleotide doublet as any two ribonucleotides that have any non-hydrogen atoms within 6.0 Å of each other. The minimal distance between the atoms of two nucleotides in a doublet was assigned as the distance between the doublet nucleotides. Then, we defined a long-range nucleotide doublet as a nucleotide doublet of two ribonucleotides that belong to distant RNA secondary structure elements, stems or loops. For that, we used the definitions from [20] (Scheme 1).

**Scheme 1.**
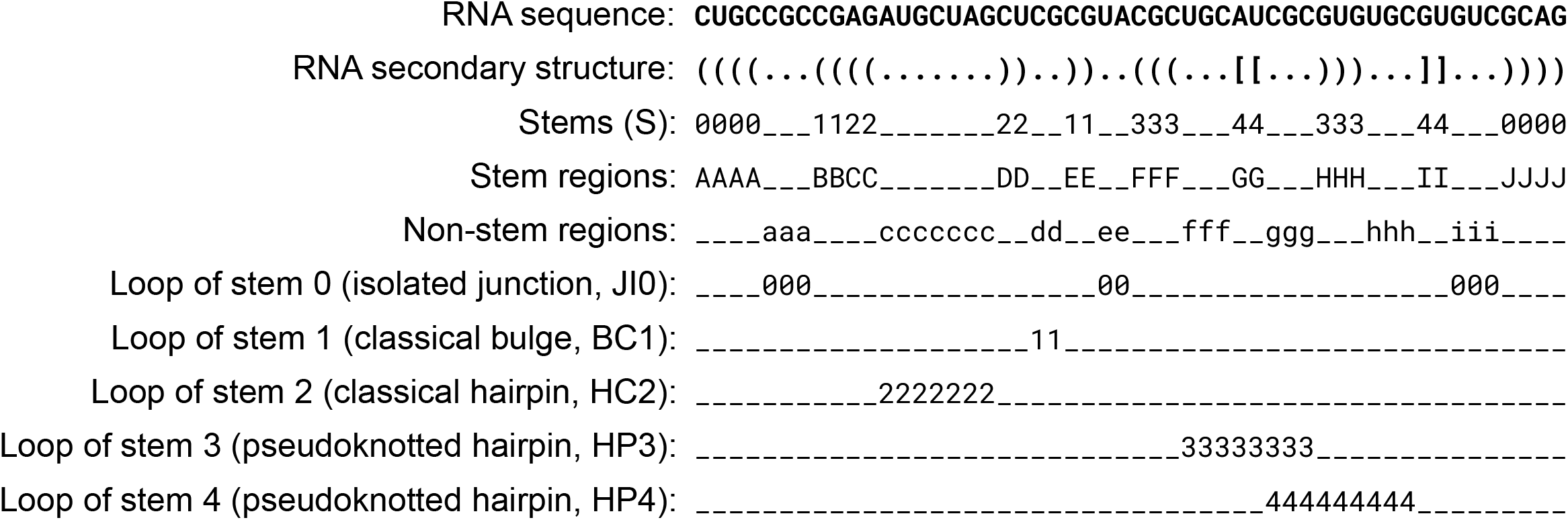
RNA secondary structure description scheme. The presented RNA sequence and structure is an artificial example designed for demonstration.

According to these definitions, a *stem* is defined as a maximal set of N ≥ 2 consecutive Watson-Crick (A-U, G-C) or Wobble (G-U) base pairs. An RNA chain is then represented as an alternating sequence of stem and non-stem regions (Scheme 1), with non-stem regions possibly being of zero length (e.g., absent non-stem region b on Scheme 1). Each stem (S) confines a set of regions that form a *loop*. Dangling ends are treated as a loop of a dummy stem that confines the entire sequence. A stem and a loop are called *adjacent* if at least a pair of their regions are neighbors in the RNA sequence. In relation to adjacent stems, each loop belongs to one of four types: a *hairpin* (H), a *bulge* (B), an *internal loop* (I), or a *multiple junction* (J). A bulge is a special case of an internal loop that involves a region of zero length. In relation to pseudoknots, each loop belongs to one of three classes: *classical* loop (C, distant from pseudoknots), *isolated* loop (I, adjacent to a pseudoknot), or *pseudoknotted* loop (P, part of a pseudoknot). Then, two RNA secondary structure elements, a stem and a loop, two stems, or two loops, are called *distant* from each other if they are not adjacent and the sets of their adjacent elements do not overlap, i.e., they do not share a third element that is adjacent to both. E.g., on Scheme 1 HC2 and S3 are distant, but HP3 and JI0 are not distant as they are both adjacent to S3. For the strict formal description of the used definitions see Supplementary Text in [20].

In case of more than one RNA chain forming together an RNA secondary structure, an intermolecular long-range nucleotide doublet was added to the dataset if at least one of its nucleotides belonged to an RNA chain from the non-redundant set of RNA structures. Overall, the formed dataset included 17,272 long-range nucleotide doublets formed by 70 RNA chains from 62 PDB entries of the non-redundant set (Table S1).

### 2. Annotation of motifs and interaction types

To annotate the long-range doublets with the commonly known motifs and interaction types we used the five conventional tools: FR3D [37], DSSR [38], MC-Annotate [39], ClaRNA [40], and RNAView [41] (Table 3). Fora number of common motifs, we also used the urslib2 python library [20]. Unfortunately, we could not use the NASSAM tool [64], as there is only a web server version available and it failed to work on some of the PDB entries (e.g., 6HIW).

**Table 3.**
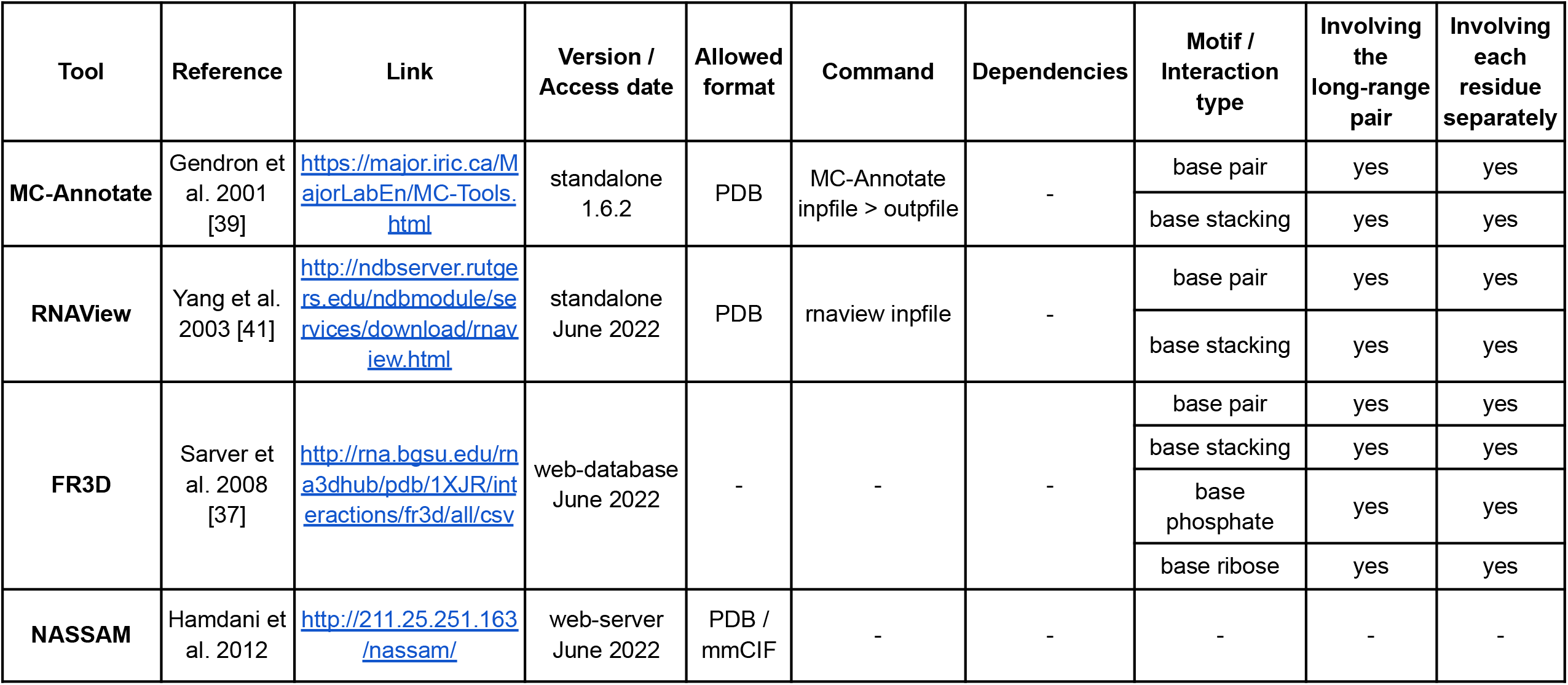
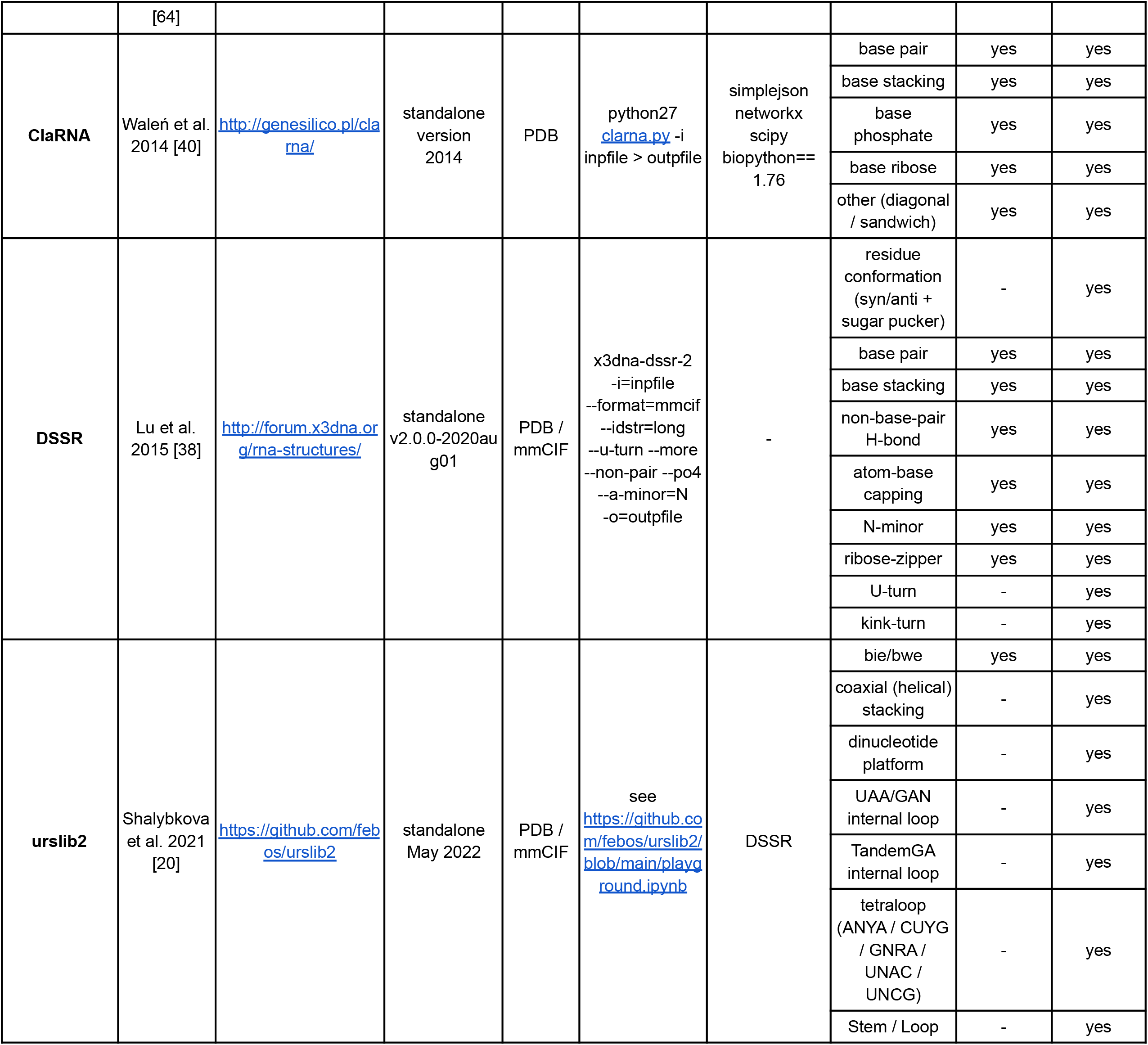
Overview of the tools and motifs and interaction types used to annotate the dataset of long-range residue-residue pairs.

Where applicable, we also annotated motifs and interaction types involving only one of the nucleotides of a doublet. In the case of FR3D, we parsed the annotations available in the RNA Structure Atlas [23] without running the tool explicitly. To run PDB-only tools on mmCIF-only PDB entries we used pdb-bundles provided by RCSB PDB [63]. We applied the default score threshold ≥ 0.5 for all ClaRNA annotations used in this work. Separately, we also stored ClaRNA base pair annotations of any score.

### 3. Assembly of long-range RNA 3D modules

To assemble long-range RNA 3D modules from the long-range nucleotide doublets we used the following approach.

Two doublets are considered to belong to the same module if (1) they share one common residue and (2) the minimal distance between the non-hydrogen atoms of the other two residues is less than an *ad hoc* threshold of 6.0 Å. The second rule prevents forming of large clumps composed of close but independent interaction networks. Then, the modules are defined as connected components of a graph with the doublets as vertices and belonging to the same module as edges.

Overall, the 17,272 long-range doublets were grouped into 1,264 long-range RNA 3D modules. The modules formed a dataset called LORA (LOng-RAnge). The LORA dataset stores modules both in PDB and mmCIF formats and is available in the form of a GitHub repository (https://github.com/febos/LORA). Each module is assigned an identifier composed of a PDB entry and an ordinal number separated by an underscore, e.g., *“7o7y_7”*.

### 4. Pairwise superposition of RNA 3D modules

To superimpose two arbitrary RNA 3D modules we developed a new algorithm called ARTEM (Aligning Rna TErtiary Motifs). The algorithm relies on an assumption that in the best possible superposition of two similar RNA 3D modules at least one residue has a counterpart with RMSD close to zero, and works as follows.

For two RNA modules X and Y of sizes N and M, for every (i, j) pair, where 1 ≤ i ≤ N and 1 ≤ j ≤ M, do the following:

1. Superimpose two modules using the Kabsch algorithm [65] considering only 5-atom representations of the residues X_i_ and Y_i_;
2. Calculate the centers of mass of 5-atom representations of all the residues of the two superimposed modules;
3. Based on the calculated centers of mass identify a subset of mutually closest residues at a distance < *MATCHRANGE* Å, such that if a residue X_s_ is the closest counterpart to a residue Y_t_, the residue Y_t_ is the closest counterpart to the residue X_s_, and the distance between their centers of mass is less than *MATCHRANGE* Å, then the pair (X_s_, Y_t_) belongs to the subset.
4. Superimpose the modules using the Kabsch algorithm considering only 3-atom base representations of the mutually closest residues subset. Calculate *RMSD* on the set of atoms used for the superposition. Output *RMSD* and *SIZE* (mutually closest subset size) values, as well as the coordinates of the superimposed module Y.

Here, module X is considered a reference, and module Y is a query, hence the module X coordinates are constant. For the 5-atom representation, we use a scheme close to the one used in SimRNA [66]: the representation includes the P atom, the center of mass of the ribose atoms, and three base atoms - N9, C2, and C6 for purines, and N1, C2, and C4 for pyrimidines. For the 3-atom base representation, the same three base atoms are used. The *MATCHRANGE* threshold was chosen manually and set to 3.0 Å to eliminate random “noise” matchings. The 3-atom representation is used in step 4 instead of the 5-atom representation to facilitate the superposition of similar base arrangements having different backbone topologies. For all the modified residues present in the used set of RNA structures we assigned 5-atom representations manually by analogy with their closest standard residues.

ARTEM is able to identify similar 3D arrangements of bases in RNA 3D structures regardless of their sequence, backbone topology, and annotated interaction types. The theoretical time complexity of ARTEM is O(n^4^), where n is the total number of atoms in the two modules, as the Kabsch algorithm of complexity O(n) [67] and the search for mutually closest residues of complexity O(n^2^) are being run O(N*M) ≈ O(n^2^) times, giving O(n+n^2^)*O(n^2^) = O(n^4^). A parallelized Python implementation of ARTEM is available in the form of a GitHub repository (https://github.com/david-bogdan-r/ARTEM). The implementation uses k-d trees [68] for the mutually closest residues search which on average works better than the calculation of all the pairwise distances running O(n^2^) in time. The tool reads and writes both PDB and mmCIF formats. The user is able to specify the particular residues of interest or run ARTEM for the entire coordinate files. Also, the user can choose to save the superimposed query structures in the preferred format. The implementation with default parameters takes around one minute to run an entire 5,970-residue eukaryotic ribosome (PDB entry 7O7Y) against a 160-residue TPP riboswitch (PDB entry 2GDI) on 32 cores, taking under 2Gb RAM at peak on an AMD Ryzen 9 5950X machine with 128Gb RAM. On the same machine on 32 cores a run of a 2,828-residue LSU rRNA (PDB entry 1FFK) against itself requires 20 minutes in time and 70Gb of RAM.

### 5. Identification of long-range RNA tertiary motifs

To identify the most common tertiary motifs among the long-range RNA 3D modules of the LORA dataset (LORA modules) we developed the following original approach.

First, we defined a *long-range tertiary motif instance* as a LORA module sub-structure (a subset of residues) that involves at least one long-range doublet and produces at least one good structural superposition with any other sub-structure of another LORA module using the ARTEM algorithm. A good structural superposition was defined as a superposition of *RMSD* and *SIZE* satisfying two conditions: *RMSD* < 3.0 Å and *RMSD/SIZE* < 0.25 Å. Thus, a long-range tertiary motif instance according to the above definition can be considered a reflection of a local structural similarity between two LORA modules. The thresholds were chosen based on the visual examination of the results from preliminary experiments. The more strict thresholds resulted in a number of false negatives, i.e., missing relevant structural similarities. In contrast, the relaxed thresholds resulted in a significant number of false positives, i.e., accidental close arrangements of centers of mass of the residues with completely different residue orientations. Therefore, the thresholds were chosen to minimize the number of false negatives and keep the number of false positives as limited as possible.

Then, we performed a motif instance search for each LORA module versus all the other LORA modules. Overall, we identified more than one million (1,156,112) long-range tertiary motif instances. Each motif instance was assigned a unique identifier made of a source module identifier and an ordinal number, e.g., *“6th6_18_261”* is an instance number 261 derived from module *6th6_18*. To derive the most frequent motifs from the motif instances of sizes from 2 to 30 we applied a heuristic two-step clustering procedure. The procedure was applied for each motif size separately and was designed to find the 100 most frequent motifs of each size and to minimize the number of computationally intensive pairwise comparisons with ARTEM. The logic behind the procedure is to find the centers of the “dense spots” of the motif instance space. The procedure works as follows:

Step 1:

1.1) Sort all the motif instances of a given size N in the decreasing order of the number of the good superpositions found;
1.2) Treat the first instance as a center of the first cluster;
1.3) For each of the remaining instances:
  1.3.1) For each currently existing cluster:
    - If the instance produces a good superposition of size N with the center of the cluster, put the instance into the cluster and go to the next instance;
  1.3.2) If no good size N superposition is found, add a new cluster to the end of the clusters list and make the current instance its center;
  1.3.3) After every 1,000 instances, sort the clusters list in the decreasing order of the number of instances and keep only the first 1,000 clusters.

As a result of step 1 we have 1,000 cluster centers that are presumably close to the instance space “dense spots”. Then, step 2 is applied for each of the 100 most populated clusters to find the proper centers of the “dense spots”.

Step 2:

2) For each cluster center:

2.1) Among all the motif instances of size N (i.e., not only cluster members) select the instances producing a good size N superposition with the current center. Treat the current center along with the selected instances as the current cluster;
2.2) Calculate all pairwise RMSD between the cluster members using 3-atom base representations and choose an instance with the least median RMSD as a new current center;
2.3) If the new center and the previous center are the same instance - output the instance as a “dense spot” center, otherwise - return to step 2.1.

The resulting “dense spot” centers are then called *motif representatives* or just *motifs*. The suggested approach has three important limitations:

1. The final clusters are not mutually exclusive, i.e., one instance can belong to several clusters.
2. Instances of different “canonically defined” tertiary motifs (e.g., type I A-minor and type II A-minor) can belong to the same cluster if a good superposition can be found between them. Yet, if a canonically defined tertiary motif is highly populated, i.e., forms a “dense spot”, it’s highly likely that it will be represented among the centers of populated clusters.
3. Given the first two limitations the resulting cluster sizes should not be treated as quantitative measures but rather qualitative ones. Thus, it’s not safe to state that a 501-instance cluster center is more populated than a 500-instance cluster center, but it can be safely treated as more populated than a 10-instance cluster center.

Considering the above, we believe the proposed approach is able to identify the most common long-range tertiary motifs characteristic of non-coding RNA 3D structures, although for their detailed quantitative analysis regarding e.g., sequence specificity additional filtering is required.

Each motif is assigned a unique identifier of the format *clustersize_motifsize_uniqueness_representative*, e.g., *“641_8_0.52_6th6_65_738”*.

Here, *clustersize* is the number of instances within the current cluster, *motifsize* is the motif size in residues, and *representative* is the identifier of the representative motif instance. *uniqueness* is a measure of the motif “originality” and is calculated as follows. A given cluster is being compared with all the more populated clusters of the same *motifsize* and for each comparison, the share of the current cluster instances not belonging to the more populated cluster is calculated, and the smallest of the shares is then assigned as the motif *uniqueness*. The *uniqueness* value provides easier navigation through the motifs helping to identify alternative centers of the same “dense spot” or centers of overlapping “dense spots”.

The LORA dataset includes the table specifying the residues of all the 1,156,112 motif instances. All the resulting motifs (i.e., cluster centers) are available in mmCIF and PDB formats. For every cluster of at least 100 instances, the dataset includes all its members in mmCIF and PDB formats along with a table of their residue matchings, RMSD, and per-residue RMSD in relation to the motif representative.

The entire two-step clustering procedure was completed in 253 hours on 32 cores on an AMD Ryzen 9 5950X machine with 128Gb RAM.

## DATA AVAILABILITY

The LORA dataset and all the supplementary materials are available at https://github.com/febos/LORA. An implementation of the ARTEM algorithm is available at https://github.com/david-bogdan-r/ARTEM.

## ACKNOWLEDGEMENTS

E.F.B. wishes to thank his former supervisors Mikhail Roytberg and Ivan Kulakovskiy, who taught him priceless principles for conducting research that led to this work. Also, E.F.B. wishes to thank his daughter Leia for a walk in the park during which he came up with the idea of the ARTEM algorithm.

## SUPPLEMENTARY MATERIALS

**Figure S1.**
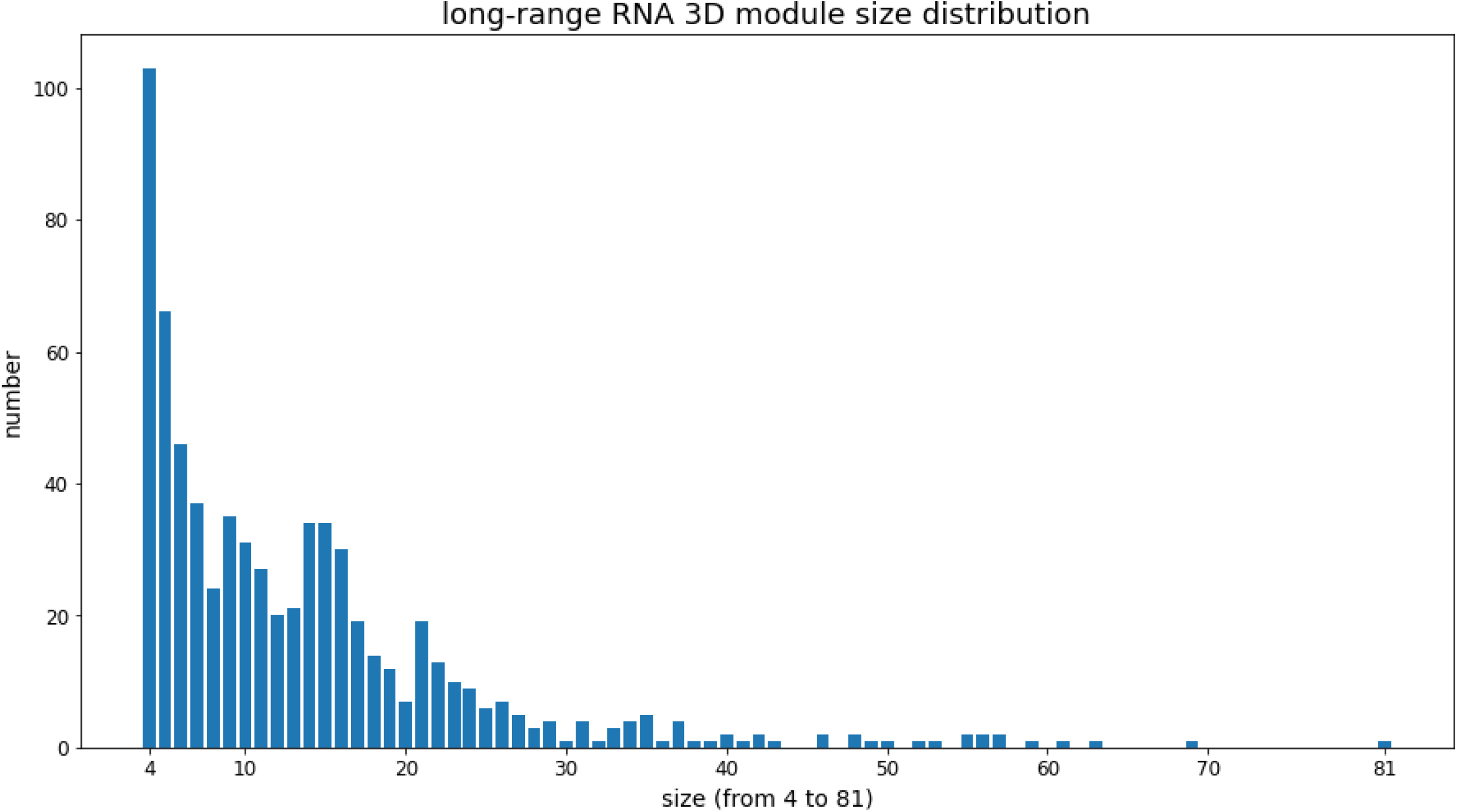
Distribution of the assembled long-range RNA 3D modules relative to their size in residues. The modules of size two, size three, and sizes higher than 100 are not shown.

**Figure S2.**
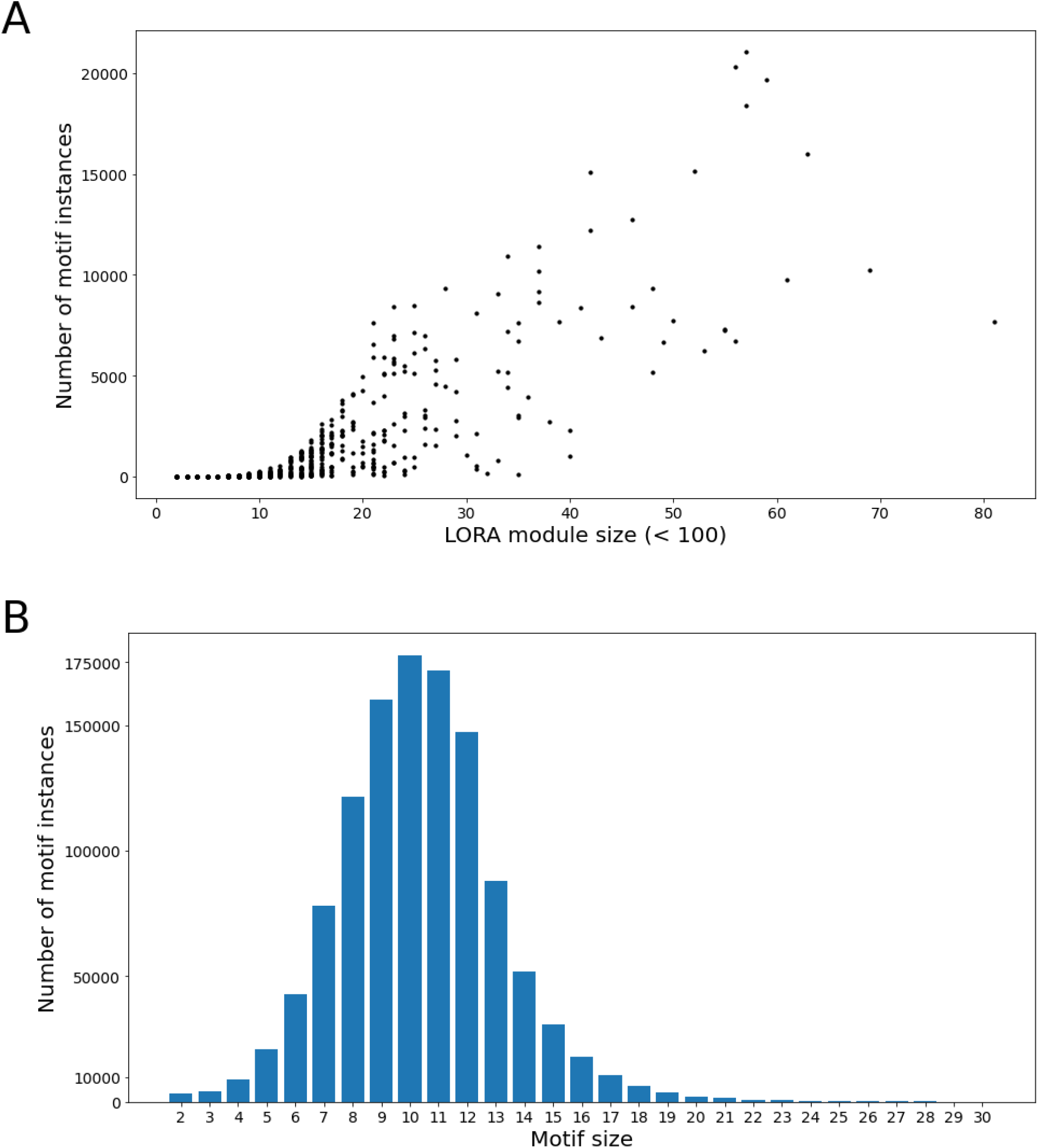
(A) Distribution of the number of motif instances produced by each LORA module relative to the module size. Each dot represents a LORA module. Modules of sizes larger than 100 residues are omitted for clarity. (B) Distribution of the number of motif instances relative to their size. Instances larger than 30 residues are omitted for clarity.

**Figure S3.**
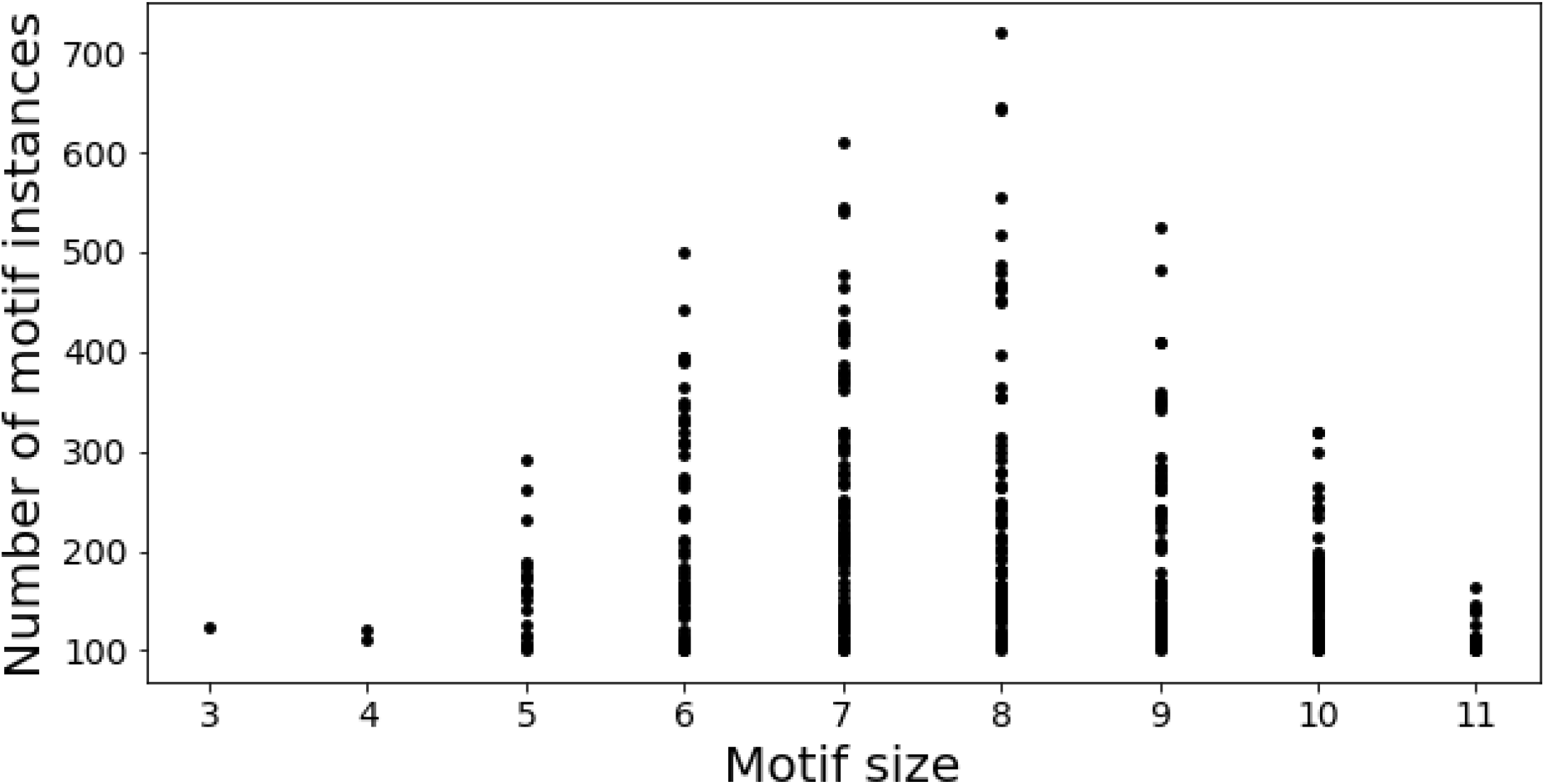
Distribution of the number of instances relative to the motif size of the 424 most populated motifs with at least 100 instances. Each dot represents a motif.

## REFERENCES

1) Santosh, B., Varshney, A., & Yadava, P. K. (2015). Non-coding RNAs: biological functions and applications. Cell biochemistry and function, 33(1), 14–22. DOI: 10.1002/cbf.3079

2) Johnsson, P., Lipovich, L., Grandér, D., & Morris, K. V. (2014). Evolutionary conservation of long non-coding RNAs; sequence, structure, function. Biochimica et Biophysica Acta (BBA)-General Subjects, 1840(3), 1063–1071. DOI: 10.1016/j.bbagen.2013.10.035

3) Leontis, N. B., Lescoute, A., & Westhof, E. (2006). The building blocks and motifs of RNA architecture. Current opinion in structural biology, 16(3), 279–287. DOI: 10.1016/j.sbi.2006.05.009

4) Hendrix, D. K., Brenner, S. E., & Holbrook, S. R. (2005). RNA structural motifs: building blocks of a modular biomolecule. Quarterly reviews of biophysics, 38(3), 221–243. DOI: 10.1017/S0033583506004215

5) Butcher, S. E., & Pyle, A. M. (2011). The molecular interactions that stabilize RNA tertiary structure: RNA motifs, patterns, and networks. Accounts of chemical research, 44(12), 1302–1311. DOI: 10.1021/ar200098t

6) Nissen, P., Ippolito, J. A., Ban, N., Moore, P. B., & Steitz, T. A. (2001). RNA tertiary interactions in the large ribosomal subunit: the A-minor motif. Proceedings of the National Academy of Sciences, 98(9), 4899–4903. DOI: 10.1073/pnas.081082398

7) Chan, C. W., Chetnani, B., & Mondragón, A. (2013). Structure and function of the T-loop structural motif in noncoding RNAs. Wiley Interdisciplinary Reviews: RNA, 4(5), 507–522. DOI: 10.1002/wrna.1175

8) Batey, R. T., Rambo, R. P., & Doudna, J. A. (1999). Tertiary motifs in RNA structure and folding. Angewandte Chemie International Edition, 38(16), 2326–2343. DOI: 10.1002/(SICI)1521-3773(19990816)38:16<2326::AID-ANIE2326>3.0.CO;2-3

9) Tamura, M., & Holbrook, S. R. (2002). Sequence and structural conservation in RNA ribose zippers. Journal of molecular biology, 320(3), 455–474. DOI: 10.1016/S0022-2836(02)00515-6

10) Zirbel, C. L., Šponer, J. E., Šponer, J., Stombaugh, J., & Leontis, N. B. (2009). Classification and energetics of the base-phosphate interactions in RNA. Nucleic acids research, 37(15), 4898–4918. DOI: 10.1093/nar/gkp468

11) Leontis, N. B., & Westhof, E. (2001). Geometric nomenclature and classification of RNA base pairs. Rna, 7(4), 499–512. DOI: 10.1017/S1355838201002515

12) Mládek, A., Sponer, J. E., Kulhánek, P., Lu, X. J., Olson, W. K., & Sponer, J. (2012). Understanding the sequence preference of recurrent RNA building blocks using quantum chemistry: the intrastrand RNA dinucleotide platform. Journal of chemical theory and computation, 8(1), 335–347. DOI: 10.1021/ct200712b

13) Baulin, E., Metelev, V., & Bogdanov, A. (2020). Base-intercalated and base-wedged stacking elements in 3D-structure of RNA and RNA–protein complexes. Nucleic acids research, 48(15), 8675–8685. DOI: 10.1093/nar/gkaa610

14) Chawla, M., Chermak, E., Zhang, Q., Bujnicki, J. M., Oliva, R., & Cavallo, L. (2017). Occurrence and stability of lone pair-π stacking interactions between ribose and nucleobases in functional RNAs. Nucleic acids research. DOI: 10.1093/nar/gkx757

15) Montemayor, E. J., Virta, J. M., Hagler, L. D., Zimmerman, S. C., & Butcher, S. E. (2019). Structure of an RNA helix with pyrimidine mismatches and cross-strand stacking. Acta Crystallographica Section F: Structural Biology Communications, 75(10), 652–656. DOI: 10.1107/S2053230X19012172

16) Geary, C., Baudrey, S., & Jaeger, L. (2008). Comprehensive features of natural and in vitro selected GNRA tetraloop-binding receptors. Nucleic acids research, 36(4), 1138–1152. DOI: 10.1093/nar/gkm1048

17) Huang, L., & Lilley, D. M. (2016). The kink turn, a key architectural element in RNA structure. Journal of molecular biology, 428(5), 790–801. DOI: 10.1016/j.jmb.2015.09.026

18) Lee, J. C., Gutell, R. R., & Russell, R. (2006). The UAA/GAN internal loop motif: a new RNA structural element that forms a cross-strand AAA stack and long-range tertiary interactions. Journal of molecular biology, 360(5), 978–988. DOI: 10.1016/j.jmb.2006.05.066

19) Steinberg, S. V., & Boutorine, Y. I. (2007). G-ribo: a new structural motif in ribosomal RNA. Rna, 13(4), 549–554. DOI: 10.1261/rna.387107

20) Shalybkova, A. A., Mikhailova, D. S., Kulakovskiy, I. V., Fakhranurova, L. I., & Baulin, E. F. (2021). Annotation of the local context of RNA secondary structure improves the classification and prediction of A-minors. RNA, 27(8), 907–919. DOI: 10.1261/rna.078535.120

21) Gianfrotta, C., Reinharz, V., Barth, D., & Denise, A. (2021). A Graph-Based Similarity Approach to Classify Recurrent Complex Motifs from Their Context in RNA Structures. 19th Symposium on Experimental Algorithms. DOI: 10.4230/LIPIcs.SEA.2021.19

22) Laing, C., Wen, D., Wang, J. T., & Schlick, T. (2012). Predicting coaxial helical stacking in RNA junctions. Nucleic acids research, 40(2), 487–498. DOI: 10.1093/nar/gkr629

23) Petrov, A. I., Zirbel, C. L., & Leontis, N. B. (2013). Automated classification of RNA 3D motifs and the RNA 3D Motif Atlas. Rna, 19(10), 1327–1340. DOI: 10.1261/rna.039438.113

24) Popenda, M., Szachniuk, M., Blazewicz, M., Wasik, S., Burke, E. K., Blazewicz, J., & Adamiak, R. W. (2010). RNA FRABASE 2.0: an advanced web-accessible database with the capacity to search the three-dimensional fragments within RNA structures. BMC bioinformatics, 11(1), 1–12. DOI: 10.1186/1471-2105-11-231

25) Richardson, K. E., Kirkpatrick, C. C., & Znosko, B. M. (2020). RNA CoSSMos 2.0: an improved searchable database of secondary structure motifs in RNA three-dimensional structures. Database, 2020. DOI: 10.1093/database/baz153

26) Bindewald, E., Hayes, R., Yingling, Y. G., Kasprzak, W., & Shapiro, B. A. (2008). RNAJunction: a database of RNA junctions and kissing loops for three-dimensional structural analysis and nanodesign. Nucleic acids research, 36(suppl_1), D392–D397. DOI: 10.1093/nar/gkm842

27) Chojnowski, G., Waleń, T., & Bujnicki, J. M. (2014). RNA Bricks—a database of RNA 3D motifs and their interactions. Nucleic acids research, 42(D1), D123–D131. DOI: 10.1093/nar/gkt1084

28) Tamura, M., Hendrix, D. K., Klosterman, P. S., Schimmelman, N. R., Brenner, S. E., & Holbrook, S. R. (2004). SCOR: Structural Classification of RNA, version 2.0. Nucleic acids research, 32(suppl_1), D182–D184. DOI: 10.1093/nar/gkh080

29) Wu, L., Chai, D., Fraser, M. E., & Zimmerly, S. (2012). Structural variation and uniformity among tetraloop-receptor interactions and other loop-helix interactions in RNA crystal structures. PLoS One, 7(11), e49225. DOI: 10.1371/journal.pone.0049225

30) Xin, Y., Laing, C., Leontis, N. B., & Schlick, T. (2008). Annotation of tertiary interactions in RNA structures reveals variations and correlations. Rna, 14(12), 2465–2477. DOI: 10.1261/rna.1249208

31) Nagaswamy, U., & Fox, G. E. (2002). Frequent occurrence of the T-loop RNA folding motif in ribosomal RNAs. Rna, 8(9), 1112–1119. DOI: 10.1017/S135583820202006X

32) Krasilnikov, A. S., & Mondragón, A. (2003). On the occurrence of the T-loop RNA folding motif in large RNA molecules. Rna, 9(6), 640–643. DOI: 10.1261/rna.2202703

33) Ulyanov, N. B., & James, T. L. (2010). RNA structural motifs that entail hydrogen bonds involving sugar–phosphate backbone atoms of RNA. New Journal of Chemistry, 34(5), 910–917. DOI: 10.1039/b9nj00754g

34) Appasamy, S. D., Hamdani, H. Y., Ramlan, E. I., & Firdaus-Raih, M. (2016). InterRNA: a database of base interactions in RNA structures. Nucleic acids research, 44(D1), D266–D271. DOI: 10.1093/nar/gkv1186

35) Reinharz, V., Soulé, A., Westhof, E., Waldispühl, J., & Denise, A. (2018). Mining for recurrent long-range interactions in RNA structures reveals embedded hierarchies in network families. Nucleic acids research, 46(8), 3841–3851. DOI: 10.1093/nar/gky197

36) Oliver, C., Mallet, V., Philippopoulos, P., Hamilton, W. L., & Waldispuhl, J. (2020). VeRNAl: Mining RNA Structures for Fuzzy Base Pairing Network Motifs. arXiv preprint arXiv:2009.00664. DOI: 10.48550/arXiv.2009.00664

37) Sarver, M., Zirbel, C. L., Stombaugh, J., Mokdad, A., & Leontis, N. B. (2008). FR3D: finding local and composite recurrent structural motifs in RNA 3D structures. Journal of mathematical biology, 56(1), 215–252. DOI: 10.1007/s00285-007-0110-x

38) Lu, X. J., Bussemaker, H. J., & Olson, W. K. (2015). DSSR: an integrated software tool for dissecting the spatial structure of RNA. Nucleic acids research, 43(21), e142–e142. DOI: 10.1093/nar/gkv716

39) Gendron, P., Lemieux, S., & Major, F. (2001). Quantitative analysis of nucleic acid three-dimensional structures. Journal of molecular biology, 308(5), 919–936. DOI: 10.1006/jmbi.2001.4626

40) Waleń, T., Chojnowski, G., Gierski, P., & Bujnicki, J. M. (2014). ClaRNA: a classifier of contacts in RNA 3D structures based on a comparative analysis of various classification schemes. Nucleic acids research, 42(19), e151–e151. DOI: 10.1093/nar/gku765

41) Yang, H., Jossinet, F., Leontis, N., Chen, L., Westbrook, J., Berman, H., & Westhof, E. (2003). Tools for the automatic identification and classification of RNA base pairs. Nucleic acids research, 31(13), 3450–3460. DOI: 10.1093/nar/gkg529

42) Ferrè, F., Ponty, Y., Lorenz, W. A., & Clote, P. (2007). DIAL: a web server for the pairwise alignment of two RNA three-dimensional structures using nucleotide, dihedral angle and base-pairing similarities. Nucleic acids research, 35(suppl_2), W659–W668. DOI: 10.1093/nar/gkm334

43) Piątkowski, P., Jabłońska, J., Żyła, A., Niedziałek, D., Matelska, D., Jankowska, E., Waleń, T., Dawson, W. K. & Bujnicki, J. M. (2017). SupeRNAlign: a new tool for flexible superposition of homologous RNA structures and inference of accurate structure-based sequence alignments. Nucleic acids research, 45(16), e150–e150. DOI: 10.1093/nar/gkx631

44) Rahrig, R. R., Leontis, N. B., & Zirbel, C. L. (2010). R3D Align: global pairwise alignment of RNA 3D structures using local superpositions. Bioinformatics, 26(21), 2689–2697. DOI: 10.1093/bioinformatics/btq506

45) Zheng, J., Xie, J., Hong, X., & Liu, S. (2019). RMalign: an RNA structural alignment tool based on a novel scoring function RMscore. BMC genomics, 20(1), 1–10. DOI: 10.1186/s12864-019-5631-3

46) Chang, Y. F., Huang, Y. L., & Lu, C. L. (2008). SARSA: a web tool for structural alignment of RNA using a structural alphabet. Nucleic acids research, 36(suppl_2), W19–W24. DOI: 10.1093/nar/gkn327

47) Szachniuk, M. (2019). RNApolis: computational platform for RNA structure analysis. Foundations of Computing and Decision Sciences, 44(2), 241–257. DOI: 10.2478/fcds-2019-0012

48) Wang, C. W., Chen, K. T., & Lu, C. L. (2010). iPARTS: an improved tool of pairwise alignment of RNA tertiary structures. Nucleic acids research, 38(suppl_2), W340–W347. DOI: 10.1093/nar/gkq483

49) Yang, C. H., Shih, C. T., Chen, K. T., Lee, P. H., Tsai, P. H., Lin, J. C., Yen, C. Y., Lin, T. Y. & Lu, C. L. (2016). iPARTS2: an improved tool for pairwise alignment of RNA tertiary structures, version 2. Nucleic acids research, 44(W1), W328–W332. DOI: 10.1093/nar/gkw412

50) Islam, S., Rahaman, M. M., & Zhang, S. (2021). RNAMotifContrast: a method to discover and visualize RNA structural motif subfamilies. Nucleic acids research, 49(11), e61–e61. DOI: 10.1093/nar/gkab131

51) Roll, J., Zirbel, C. L., Sweeney, B., Petrov, A. I., & Leontis, N. (2016). JAR3D Webserver: Scoring and aligning RNA loop sequences to known 3D motifs. Nucleic acids research, 44(W1), W320–W327. DOI: 10.1093/nar/gkw453

52) Duarte, C. M., Wadley, L. M., & Pyle, A. M. (2003). RNA structure comparison, motif search and discovery using a reduced representation of RNA conformational space. Nucleic Acids Research, 31(16), 4755–4761. DOI: 10.1093/nar/gkg682

53) Chen, X., Khan, N. S., & Zhang, S. (2020). LocalSTAR3D: a local stack-based RNA 3D structural alignment tool. Nucleic acids research, 48(13), e77–e77. DOI: 10.1093/nar/gkaa453

54) Čech, P., Svozil, D., & Hoksza, D. (2012). SETTER: web server for RNA structure comparison. Nucleic acids research, 40(W1), W42–W48. DOI: 10.1093/nar/gks560

55) Nguyen, M. N., Sim, A. Y., Wan, Y., Madhusudhan, M. S., & Verma, C. (2017). Topology independent comparison of RNA 3D structures using the CLICK algorithm. Nucleic acids research, 45(1), e5–e5. DOI: 10.1093/nar/gkw819

56) Doherty, E. A., Batey, R. T., Masquida, B., & Doudna, J. A. (2001). A universal mode of helix packing in RNA. Nature structural biology, 8(4), 339–343. DOI: 10.1038/86221

57) Kalvari, I., Nawrocki, E. P., Ontiveros-Palacios, N., Argasinska, J., Lamkiewicz, K., Marz, M., … & Petrov, A. I. (2021). Rfam 14: expanded coverage of metagenomic, viral and microRNA families. Nucleic Acids Research, 49(D1), D192–D200. DOI: 10.1093/nar/gkaa1047

58) Leontis, N. B., & Zirbel, C. L. (2012). Nonredundant 3D structure datasets for RNA knowledge extraction and benchmarking. In RNA 3D structure analysis and prediction (pp. 281–298). Springer, Berlin, Heidelberg. DOI: 10.1007/978-3-642-25740-7_13

59) Boccaletto, P., Stefaniak, F., Ray, A., Cappannini, A., Mukherjee, S., Purta, E., … & Bujnicki, J. M. (2022). MODOMICS: a database of RNA modification pathways. 2021 update. Nucleic Acids Research, 50(D1), D231–D235. DOI: 10.1093/nar/gkab1083

60) Sykes, M. T., & Levitt, M. (2005). Describing RNA structure by libraries of clustered nucleotide doublets. Journal of molecular biology, 351(1), 26–38. DOI: 10.1016/j.jmb.2005.06.024

61) Wang, J., Daldrop, P., Huang, L., & Lilley, D. M. (2014). The k-junction motif in RNA structure. Nucleic acids research, 42(8), 5322–5331.DOI: 10.1093/nar/gku144

62) Chetverina, H. V., Demidenko, A. A., Ugarov, V. I., & Chetverin, A. B. (1999). Spontaneous rearrangements in RNA sequences. FEBS letters, 450(1-2), 89–94. DOI: 10.1016/S0014-5793(99)00469-X

63) Burley, S. K., Berman, H. M., Bhikadiya, C., Bi, C., Chen, L., Di Costanzo, L., … & Zardecki, C. (2019). RCSB Protein Data Bank: biological macromolecular structures enabling research and education in fundamental biology, biomedicine, biotechnology and energy. Nucleic acids research, 47(D1), D464–D474. DOI: 10.1093/nar/gky1004

64) Hamdani, H. Y., Appasamy, S. D., Willett, P., Artymiuk, P. J., & Firdaus-Raih, M. (2012). NASSAM: a server to search for and annotate tertiary interactions and motifs in three-dimensional structures of complex RNA molecules. Nucleic acids research, 40(W1), W35–W41. DOI: 10.1093/nar/gks513

65) Kabsch, W. (1976). A solution for the best rotation to relate two sets of vectors. Acta Crystallographica Section A: Crystal Physics, Diffraction, Theoretical and General Crystallography, 32(5), 922–923. DOI: 10.1107/S0567739476001873

66) Boniecki, M. J., Lach, G., Dawson, W. K., Tomala, K., Lukasz, P., Soltysinski, T., Rother, K. M. & Bujnicki, J. M. (2016). SimRNA: a coarse-grained method for RNA folding simulations and 3D structure prediction. Nucleic acids research, 44(7), e63–e63. DOI: 10.1093/nar/gkv1479

67) Dolezal, R., Fronckova, K., Kirimtat, A., & Krejcar, O. (2020, June). Computational Complexity of Kabsch and Quaternion Based Algorithms for Molecular Superimposition in Computational Chemistry. In International Conference on Engineering Applications of Neural Networks (pp. 473–486). Springer, Cham. DOI: 10.1007/978-3-030-48791-1_37

68) Bentley, J. L. (1975). Multidimensional binary search trees used for associative searching. Communications of the ACM, 18(9), 509–517. DOI: 10.1145/361002.361007

